# Dissociable Anterior and Posterior Beta-Burst Dynamics Track Thought Disorder in Schizophrenia

**DOI:** 10.64898/2026.07.27.740959

**Authors:** Hsi T. Wei, Lindsey Power, Rukun Dou, Sylvain Baillet, Lena Palaniyappan

## Abstract

**Background:** Thought disorder in schizophrenia reflects disrupted internal model maintenance in predictive processing. Beta oscillations have been proposed to index top-down predictive signalling, but prior resting-state studies have focused on spectral power, overlooking transient burst dynamics that enable the stabilization and reinstatement of internal representations, especially in the default mode network (DMN).

**Methods:** Resting-state MEG was acquired from 25 patients (7 females) with schizophrenia and 25 matched controls (8 females) between May 2024 and October 2025. Beta bursts (15-30 Hz) were extracted from source-localized cortical time series, and burst rate, duration, power, and inter-regional synchrony were quantified. Associations with computationally-derived semantic and syntactic speech features were assessed using principal-component regression within patients.

**Results:** Conventional beta power did not differ between groups after correction. In contrast, beta-burst dynamics showed significant group-by-anterior–posterior-position interactions for burst rate (p = 0.018), power (p = 0.0014) and duration (p < 0.001). Within the DMN, patients showed shorter inferior parietal burst duration (t(46.0) = −2.74, FDR-p = 0.0343) and increased burst synchrony within left prefrontal cortex (t(31.0) = 2.81, FDR-p = 0.0343) and between prefrontal cortices (t(29.8) = 2.73, FDR-p = 0.0423). Within patients, burst components predicted speech organization (R² = 0.47, F(3,19) = 5.66, p = 0.006), driven by the inferior parietal burst-duration component (β = −0.55, p = 0.0036), but did not predict overall clinical severity.

**Limitations:** The modest patient sample, short resting-state recording, cross-sectional design, and lack of laminar specificity limit causal and mechanistic inference; burst-speech associations require replication.

**Conclusion:** Schizophrenia was associated with dissociable anterior and posterior beta-burst abnormalities that tracked objective speech markers of thought disorder. These findings suggest impaired hierarchical coordination of predictive beta signalling as a candidate mechanism linking intrinsic network dysfunction to disorganized thought.

## 1. Introduction

Thought disorder, clinically expressed through disorganized speech, is among the most disabling features of schizophrenia and predicts long-term functional decline ^1,2^ and treatment resistance ^3,4^. Contemporary accounts link thought disorder to impaired internal models that maintain and flexibly update internal representations in predictive processing, disrupting the coherence of self-generated thought and language (^5^ see also ^6,7^).

The brain generates internal models through the interplay of top-down predictions from anterior cortex and bottom-up sensory evidence from posterior regions ^8^, while resting-state activity helps maintain these representational hierarchies between task demands ^9^. The default mode network (DMN) is a relevant substrate for this process, particularly for self-generated and internally oriented cognition ^10,11^. However, the DMN is not unitary: posterior and anterior components differ in their coupling to perceptible events and self-generated content ^12,13^. Across this posterior-to-anterior axis, internal representation becomes progressively more abstract, from sensory-motor through lexical-semantic to self-generated narrative. DMN dynamics are consistently disrupted in schizophrenia, with connectivity and activation abnormalities linked to symptom burden ^3,14–16^.

Beta-frequency (15-30 Hz) activity is a candidate mechanism for implementing internal models. Beta activity has been proposed as a cortical signature of top-down, endogenous representation ^17^, mediating feedback from deep cortical layers to suppress conflicting input and coordinate predictions hierarchically ^18^. Rather than being sustained, beta-band activity is often organized into transient high-amplitude events ^19–21^, providing discrete windows during which cortical representations may be stabilized or reinstated (i.e., “distinct computational primitives”) ^22^. If beta bursts implement internal models, the temporal coordination of burst events across distributed regions is crucial for hierarchical integration of representations. Time-averaged spectral power can obscure this event structure ^23^, whereas burst rate, duration, peak power, and inter-regional burst coincidence (synchrony) may capture distinct aspects of beta organization ^24,25^.

Task-based evidence indicates that beta burst dynamics are disrupted in schizophrenia. During task performance, patients show reduced burst rate alongside greater burst-related BOLD activation, with more severe disorganization associated with more extensive spatial distribution of beta bursts ^26^. Resting-state studies, however, have largely focused on spectral power and temporal correlations, commonly reporting elevated frontal and reduced posterior beta power without characterizing temporal coordination of burst dynamics i.e., duration and inter-regional synchrony (Table S1). Because beta bursts may stabilize and coordinate internal models across cortical hierarchies, resting-state burst organization may directly index the integrity of predictive architecture during task-free states. Within the DMN, the antero-posterior PFC-IPL axis is the specific substrate for this: PFC generates higher level predictions; the IPL integrates contextual evidence in support of semantic concepts ^27–29^. Impaired coordination on this axis via excess synchrony (locking of top-down predictions) or as temporal instability (failure to sustain representations long enough to integrate them) may produce the conditions for predictive processing failure. Language being one of the quintessential functions of hierarchical predictive processing ^5,30^, any bias towards context-insensitive predictions, may provide the necessary conditions for thought disorder manifested in speech to emerge ^31–35^.

We examined beta burst dynamics using resting-state MEG from source-localized cortical and DMN time series, with burst rate, duration, power, and synchrony quantified using established methods ^36^. We first tested for cortex-wide spatial organization of beta-burst abnormalities, then focused on DMN subregions. This two-stage approach separates a broad cortical anterior-posterior gradient from its expression within a network directly relevant to the dynamics of spontaneous thought. Based on prior evidence of elevated frontal and reduced posterior beta power (Table S1), and impaired flexible maintenance of internal representation in schizophrenia ^37–40^, we predicted enhanced burst synchrony in anterior DMN regions and reduced temporal persistence of bursts in the IPL where semantic integration is implemented. We further predicted that these burst abnormalities would relate to objective measures of thought disorder manifested as speech disorganization ^1,41,42^.

Three findings emerge. Cortex-wide, schizophrenia was associated with a disrupted anterior-posterior gradient of burst dynamics, with enhanced frontal and attenuated posterior burst features, and no group difference in average beta power. Within the DMN, this gradient resolved to two notable effects: increased burst synchrony within and between prefrontal regions, and reduced burst duration in the inferior parietal lobule. In patients, these alterations showed dissociable behavioural correlates: burst features, especially the inferior parietal burst duration, predicted speech-organization but not the overall illness burden. The findings identify excessive prefrontal burst synchrony and reduced posterior burst persistence as a candidate spatiotemporal signature of impaired hierarchical coordination underlying disorganised speech in schizophrenia.

## 2. Methods and Materials

### 2.1 Participants

This case-control study recruited 25 community-living patients with clinically stable schizophrenia and 25 age- and sex-matched healthy controls. The study was approved by the Research Ethics Board of Douglas Mental Health University Institute (McGill University), and all participants provided written informed consent. Inclusion criteria were age 18-65 and fluency in English or French. Patients were recruited from McGill-affiliated psychiatric services and met DSM-5 criteria for schizophrenia or schizoaffective disorder. Participants completed clinical, speech, MEG, and MRI assessments. Two patients were excluded from MEG analyses due to artifacts, resulting in a final analysis sample of 48 participants (Table 1). Clinical measures included SOFAS, PANSS-8, and CGI-S. Medication load was quantified using defined daily dose (DDD) based on the ATC/DDD Index.

**Table 1.**
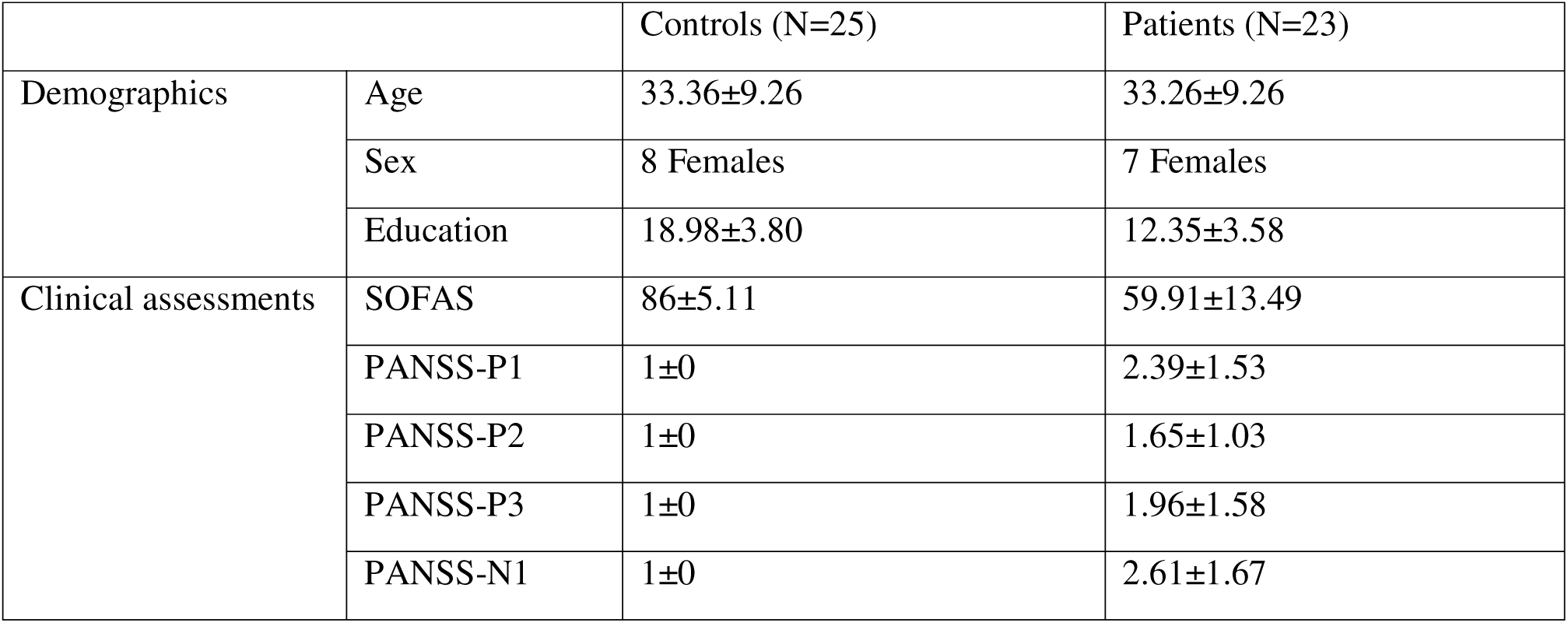
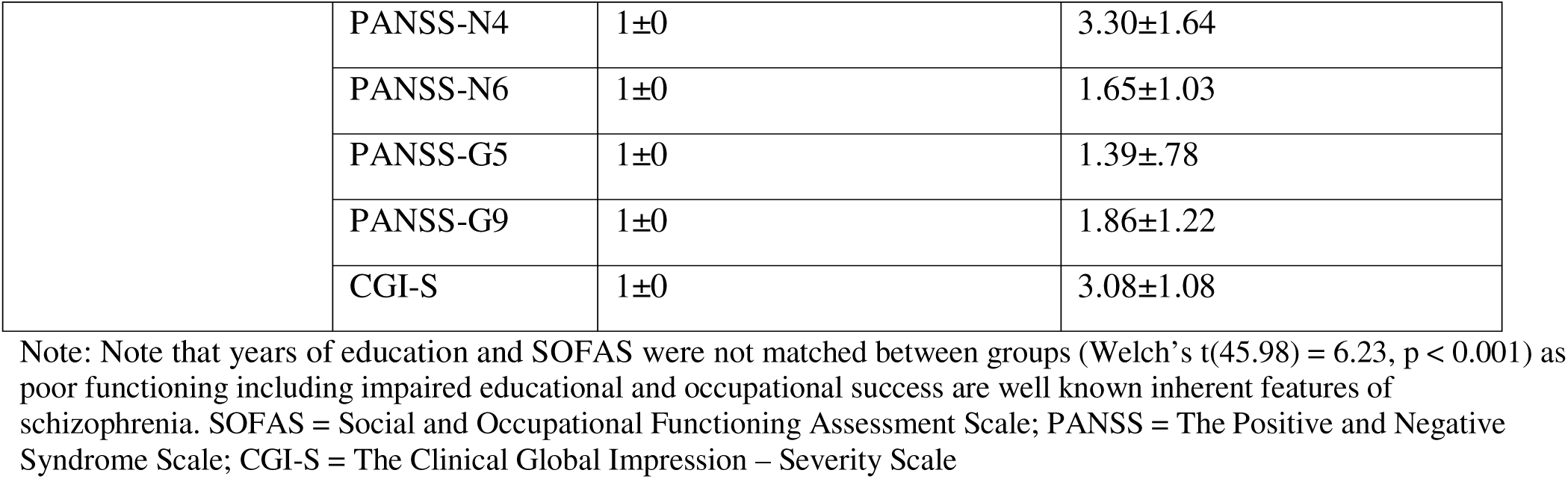
Demographics and clinical symptom scores.

|  |  | Controls (N=25) | Patients (N=23) |
| --- | --- | --- | --- |
| Demographics | Age | 33.36±9.26 | 33.26±9.26 |
|  | Sex | 8 Females | 7 Females |
|  | Education | 18.98±3.80 | 12.35±3.58 |
| Clinical assessments | SOFAS | 86±5.11 | 59.91±13.49 |
|  | PANSS-P1 | 1±0 | 2.39±1.53 |
|  | PANSS-P2 | 1±0 | 1.65±1.03 |
|  | PANSS-P3 | 1±0 | 1.96±1.58 |
|  | PANSS-N1 | 1±0 | 2.61±1.67 |
|  | PANSS-N4 | 1±0 | 3.30±1.64 |
|  | PANSS-N6 | 1±0 | 1.65±1.03 |
|  | PANSS-G5 | 1±0 | 1.39±.78 |
|  | PANSS-G9 | 1±0 | 1.86±1.22 |
|  | CGI-S | 1±0 | 3.08±1.08 |

Note: Note that years of education and SOFAS were not matched between groups (Welch's $t(45.98) = 6.23, p < 0.001$ ) as poor functioning including impaired educational and occupational success are well known inherent features of schizophrenia. SOFAS = Social and Occupational Functioning Assessment Scale; PANSS = The Positive and Negative Syndrome Scale; CGI-S = The Clinical Global Impression – Severity Scale

### 2.2 Natural speech recording and analysis

Speech samples were collected using the DISCOURSE protocol ^43^, comprising structured and free speech tasks (∼20 minutes). Recordings were transcribed using WhisperX and manually corrected, diarized for retaining only participant speech. Total word count did not differ significantly between groups (*t*(45.98) = .56, *p* = .58). Six semantic and syntactic speech features were extracted using NLP and LLM-based methods (Supplementary 1; Table S2). See Table S3 for speech feature interpretation.

### 2.3 MEG resting-state and structural MRI acquisition

Resting-state MEG data (3 minutes, eyes open) were acquired using a 275-channel CTF system at 1200 Hz. Head position was continuously monitored using fiducial coils, and EOG/ECG were recorded for artifact detection. Structural MRI was acquired on a 3T Siemens Prisma scanner for source reconstruction. Head shape was digitized using FASTRACK (Polhemus, USA; ∼100 scalp points including fiducials), and MEG-MRI co-registration was performed in Brainstorm.

### 2.4 MEG data preprocessing

Data were processed in Brainstorm using automated MATLAB pipelines. Signals were filtered at 0.1–100 Hz with a 60 Hz notch, and cardiac and ocular artifacts were removed using signal-space projection (SSP). Source reconstruction used an LCMV beamformer with unconstrained dipole orientation and overlapping spheres head models. Whole-brain parcellation used the Schaefer-200 atlas ^44^. Regional time series were extracted by reducing the three dipole orientations at each vertex to a single time course using principal component analysis (PCA), followed by sign-flipped averaging within each ROI. Analyses were restricted to the first 180 seconds to standardize data length across participants.

### 2.5 Beta oscillatory power analysis

Power spectral density was computed using Welch’s method with 4 s windows and 50% overlap. Beta power was defined as mean power in the 15-30 Hz range for each region.

### 2.6 Beta-burst analysis

Beta bursts were identified following established methods ^21^. Data were downsampled to 600 Hz, and time-frequency representations (8-35 Hz, 1 Hz resolution) were computed using Morlet wavelets (width = 10). Local maxima were identified using the Python equivalent of MATLAB’s “*imregionalmax*”. Beta bursts were defined as local maxima between 15 and 30 Hz exceeding 4× the median power, computed separately for each subject, region, and frequency. Burst onset and offset were defined using the full width at half maximum around the peak. Burst duration, normalized peak power, and burst rate were then computed.

### 2.7 Inter-regional synchrony and connectivity

Inter-regional burst synchrony within the DMN was quantified as the probability of coincident bursting between region pairs, using 150 ms epochs centred on burst onset in one region and computing burst probability in the paired region at time zero. Further details are in Supplementary 2.

Burst synchrony captures temporal co-occurrence of bursts not limited to zero-lag overlap. To validate and interpret the burst synchrony findings, beta-band amplitude envelope correlation (AEC) was computed for DMN regions (MNE-Python; bandpass-filtered at 15-30Hz using a zero-phase finite impulse response filter with a Hamming window; amplitude envelopes extracted using the Hilbert transform). Two forms of AEC were computed to distinguish between instantaneous and delayed coupling. (1) Non-orthogonalized AEC, as a continuous-signal comparator, captures the full amplitude envelope correlation between regions, including zero-lag co-fluctuations that may reflect genuine instantaneous interactions or spurious connectivity from source leakage. (2) Symmetric pairwise orthogonalized AEC removes all zero-phase-lag components from each pair of signals before computing correlations, isolating time-delayed coupling while suppressing true instantaneous interactions as well as volume conduction ^45^. Comparing results across both measures therefore identifies whether a group effect is driven primarily by instantaneous co-occurrence (including potential leakage contributions) or by sustained delayed coupling that is robust to volume conduction confounds.

Regional AEC comparisons were restricted to region pairs where burst dynamics (synchrony or duration) showed FDR-corrected group differences. To characterize overall spatial correspondence between burst synchrony group differences and each AEC type across all DMN region pairs, Mantel tests were conducted. Sliding-window AEC was computed using window lengths of 0.5–100 s, to test whether connectivity differences depended on the temporal scale of observation. Pairwise Euclidean distance between regions was computed to test whether burst synchrony differences scaled with spatial proximity, as would be expected under a signal leakage account.

### 2.8 Statistical analysis

#### 2.8.1 Group differences

Beta power and burst characteristics were compared between groups for each regional time series using independent-samples t-tests. FDR correction was applied separately for each beta characteristic across the 200 regions. Anterior-posterior (A-P) trends were assessed using linear mixed-effects models: burst_feature ∼ position*group + (position|subject), where position was defined as the mean y-coordinate of each Schaefer region.

DMN analyses grouped regions into four functionally and anatomically defined sets (Figure 2): prefrontal cortex, temporal lobe, inferior parietal lobe, and posterior cingulate cortex. Burst features were averaged within groups and compared using FDR-corrected t-tests, with correction applied separately for each burst feature across the four regional comparisons. DMN burst synchrony matrices were compared between groups, and regional synchrony was summarized within and between hemispheres. Mantel tests assessed correspondence between synchrony, connectivity, and distance matrices, with analyses repeated for AEC metrics.

Results with p < 0.05 that did not survive FDR correction are reported as trending group differences. Associations with DDD were tested to assess medication effects. For burst characteristics correlated with education, group effects adjusted for education were reported.

#### 2.8.2 Dimension reduction and regression

As pooling diagnostically distinct groups can induce spurious brain–behaviour associations, burst–speech relationships were modelled within the patient group alone (n = 23). The six burst features showing significant group differences were z-scored and summarised by their first three principal components (cumulative 85% of variance: PC1 36%, PC2 32%, PC3 16%), and the six semantic/syntactic speech features by their first principal component to provide a single speech organization factor (50% of variance). Speech organization was regressed on the three burst PCs in a single multiple linear regression; standardized coefficients (β), per-predictor significance, and the omnibus model fit are reported, with leave-one-out sensitivity to influential cases. To test symptom specificity to disorganized speech, the clinical severity component (first PC of 10 clinical scores: PANSS items, CGI-S and SOFAS; 58% of illness-burden variance; see Table S4) was regressed on the same three burst PCs. As a complementary multivariate analysis, a partial least squares (PLS) canonical correlation of the full burst and speech feature sets separately within patient and control groups, and covariate-adjusted (group, age, sex) whole sample models with 5,000-permutation are reported in the Supplement (Supplementary 3; Figure S7-9). Defined daily dose was tested against the clinical component to assess medication effects.

## 3. Results

### 3.1 Spatial gradient of burst alterations

Patients and controls did not differ in regional beta-band PSD after correction for multiple comparisons (p > 0.05). For beta burst dynamics, all three burst features showed a significant reversal of the A-P gradient between patients and controls. In healthy controls, burst power (*coef* = -4.63, *p* = 2.38e-5) and duration (*coef* = -0.0592, *p* = 3.56e-5) were greatest in posterior regions and declined toward the frontal cortex (*coef* = 0.275, *p* = 0.0143). In patients, this pattern was inverted: burst power and duration were elevated frontally and reduced posteriorly, while burst rate was disproportionately increased anteriorly (Figure 1). This group-by-position interaction was significant for burst rate (p = 0.018, coef = 0.374 ± 0.152), burst power (p = 0.0014, coef = 4.51 ± 1.32), and burst duration (p < 0.001, coef = 0.063 ± 0.016); the crossing trajectories are illustrated in Figure 1D–F. Region-wise comparisons are provided in Figure S1.

**Figure 1.**
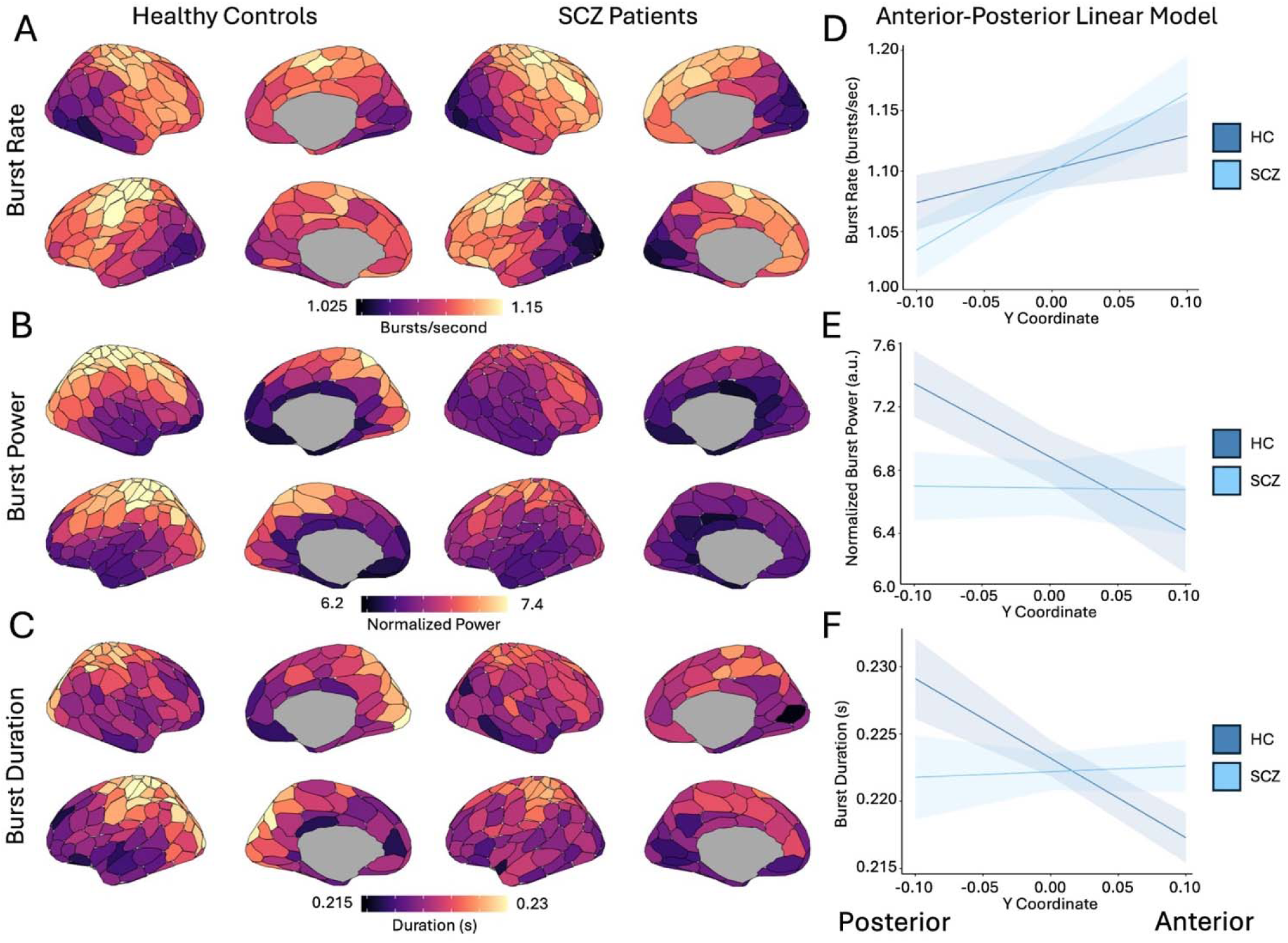
Whole-brain beta burst patterns in schizophrenia and healthy controls. (A-C) Cortical maps depicting mean burst rate (A), burst power (B), and burst duration (C) for healthy controls (left) and schizophrenia patients (right). (D-F) Linear models relating A-P position (y coordinate) to burst rate (D), burst power (E), and burst duration (F). Models are plotted separately for healthy controls (dark blue) and patients (light blue). See Supplementary Figure S1 for whole-brain beta burst characteristic summary.

Antipsychotic exposure (DDD) did not correlate with any A-P gradient within patients (all burst features r < 0.09, all p > 0.69), arguing against a medication-driven account of the spatial reversal.

### 3.2 DMN burst alterations

The cortex-wide A-P gradient was also reflected in DMN subregions. Within the DMN, patients showed shorter burst duration in the IPL, a posterior DMN node (*t*(46.0) = -2.74, *p* = 0.00858, FDR-p = 0.0343, *d* = -0.790) (Figure 2).

**Figure 2.**
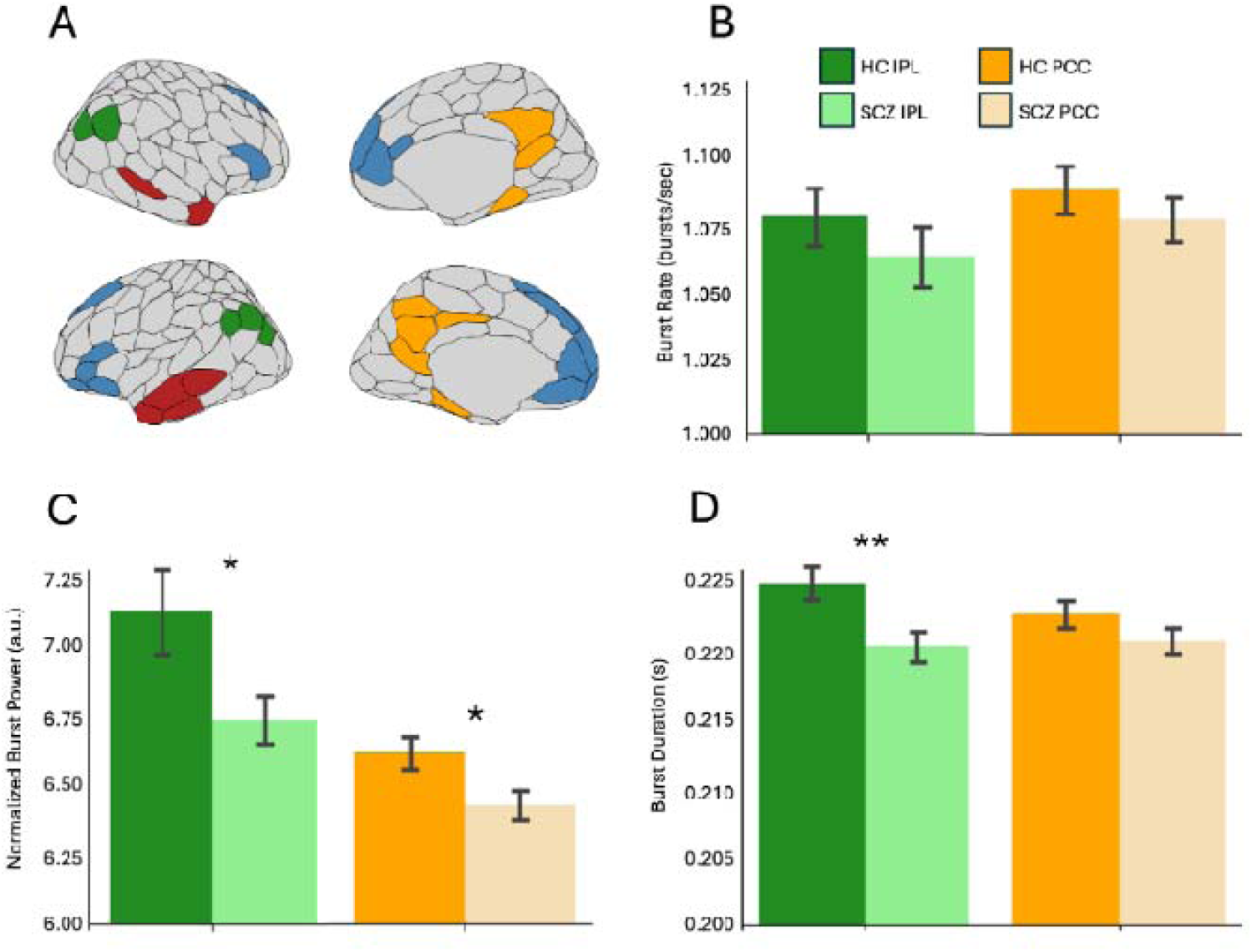
DMN beta burst characteristics in schizophrenia and healthy controls. (A) A parcellated cortical map illustrating the four DMN sub-regions extracted from the Schaefer-200 atlas: PFC (blue), temporal (red), IPL (green), and PCC (orange). (B-D) Bar graphs depicting mean IPL (green) and PCC (orange) burst rate (B), burst power (C) and burst duration (D) for healthy controls (dark) and patients (light). Error bars indicate standard error of the mean. Asterisks denote statistical significance (*: p<0.05, **: FDR-p<0.05). Bar graphs for other regions can be found in Figure S2.

There was a significantly increased zero-lag burst coincidence within left PFC (*t*(31.0) = 2.81, *p* = 0.00858, *FDR-p* = 0.0343, *d* = 0.833) and between left and right PFCs (*t*(29.8) = 2.73, *p* = 0.0106, *FDR-p* = 0.0423, *d* = 0.811) (Figure 3).

**Figure 3.**
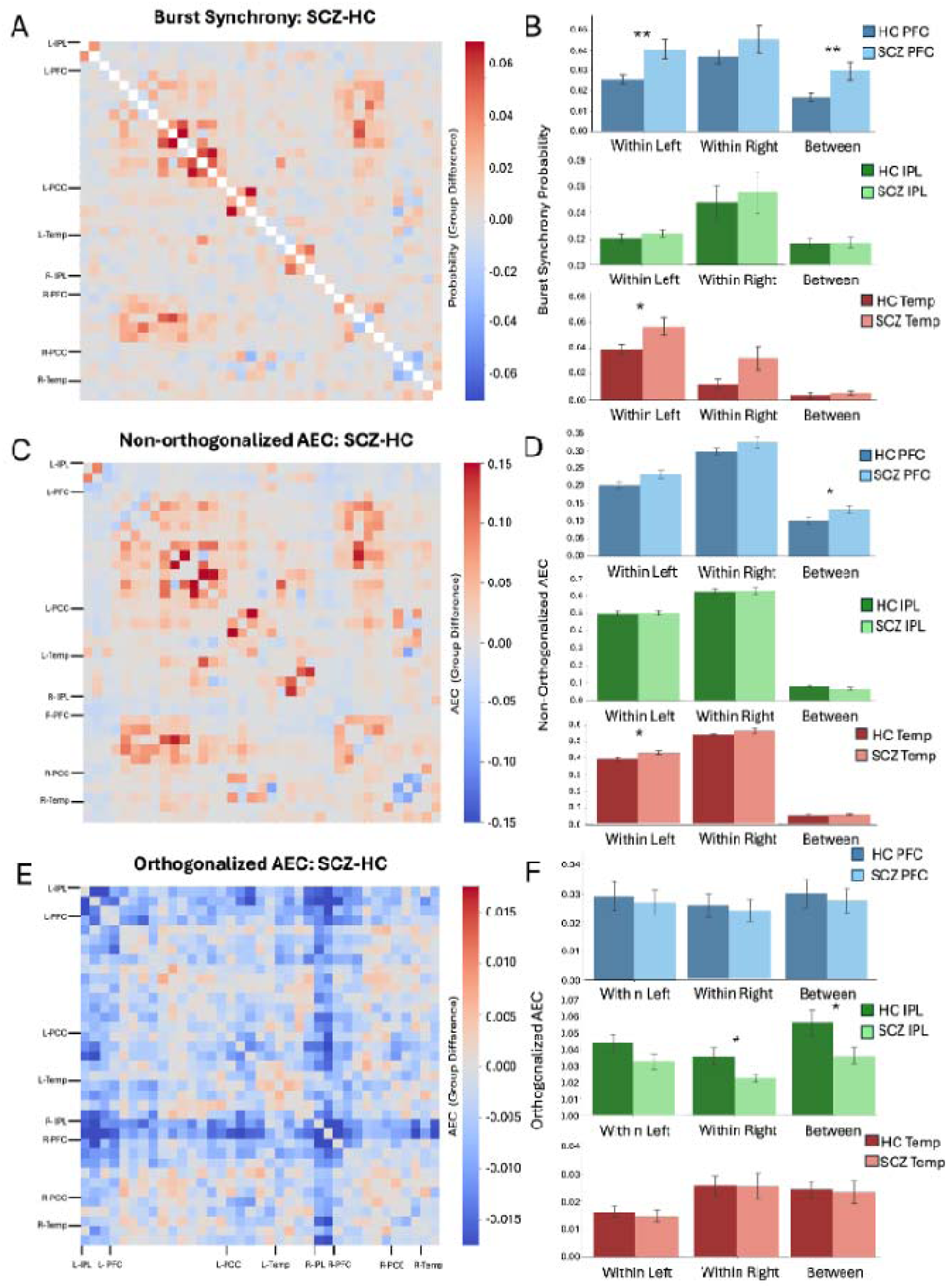
Beta burst synchrony and connectivity metrics in DMN regions. (A-B) Burst synchrony within DMN regions. (A) Group difference in synchrony (SCZ-HC) for each pair of regions in the DMN. (B; top) Mean synchrony within left PFC (left), within right PFC (center) and between left and right PFC (right) regions for healthy controls (dark blue) and patients (light blue). (B; middle) Mean synchrony within left IPL (left), with n right IPL (center) and between left and right IPL (right) regions for healthy controls (dark green) and patients (light green). (B; bottom) Mean synchrony within left temporal (left), within right temporal (center) and between left and right temporal (right) regions for healthy controls (dark red) and patients (light red). Error bars indicate standard error of the mean. Asterisks denote statistical significance (*: p<0.05, **: FDR-p<0.05). (C-D) Non-orthogonalized amplitude-envelope correlations within DMN regions. (E-F) Amplitude-envelope correlations with pairwise orthogonalization within DMN regions. Panels C-D and E-F follow the format described for panels A-B.

Increased PFC synchrony and reduced IPL burst duration were weakly negatively correlated (IPL duration with within-PFC synchrony: *r* = -0.15; with between-PFC synchrony: *r* = -0.076; Figure S3A, indicating that these two patterns may not occur in the same individuals. Within patients, DDD was not correlated with PFC synchrony (within left: *r* = -0.04, *p* = .84; between hemispheres: *r* = .17, *p* = .44) or IPL burst duration (*r* = -0.06, *p* = .80).

No significant differences in burst characteristics were observed in PFC or temporal regions of the DMN; no synchrony differences were observed within or between IPL or PCC regions (Figure S4).

Some reduction was also noted in burst power at the IPL (*t*(37.7) = -2.25, *p* = 0.0304, *d* = -0.635) and PCC (*t*(46.0) = -2.41, *p* = 0.0200, *d* = -0.694), with some increase in synchrony within the left temporal regions (*t*(36.9) = 2.19, *p* = 0.0347, *d* = 0.645) but none of these three effects reached significance after FDR correction (FDR-p > 0.06).

### 3.3 Burst synchrony and amplitude envelope connectivity

Group differences in burst synchrony across all DMN region pairs closely matched group differences in non-orthogonalized AEC (Mantel *r* = 0.857, *p* < 0.0001), confirming that the burst synchrony pattern reflects a genuine property of amplitude covariation rather than an artefact of the burst-detection approach. Burst synchrony group differences showed no spatial correspondence with orthogonalized AEC differences (Mantel *r* = 0.102, *p* = 0.076), indicating that the synchrony effect is driven primarily by zero-lag co-occurrence rather than sustained delayed coupling.

At the PFC region pairs where burst synchrony showed FDR-corrected group differences, non-orthogonalized AEC was also elevated in patients: within left PFC (*t*(30.98) = 2.81, *p* = 0.0086, *d* = 0.82) and between left and right PFC (*t*(38.8) = 2.07, *p* = 0.045, *d* = 0.61), Orthogonalized AEC showed no corresponding PFC increase, confirming that the prefrontal effect reflects instantaneous burst co-occurrence rather than sustained delayed coupling. The distance analysis further argued against a leakage account: burst synchrony differences were positively correlated with inter-regional Euclidean distance (*r* = 0.234, *p* < 0.0001; Figure S5), the opposite of what proximity-driven leakage predicts.

In contrast, at the IPL region pairs where burst duration showed an FDR-corrected group difference, orthogonalized AEC was reduced in patients (within right IPL: *t*(33.4) = -2.22, *p* = 0.034, *d* = -0.622; between left and right IPL: *t*(40.8) = -2.18, *p* = 0.035, *d* = -0.619). No corresponding IPL effect was observed for non-orthogonalized AEC.

Sliding-window AEC showed that this orthogonalized AEC reduction emerged only at window lengths of 50 s or longer, whereas non-orthogonalized AEC increases in PFC were consistent across all timescales (0.5–100 s; Figure 4). Reduced IPL coupling was associated with shorter IPL burst duration (*r* = 0.41, *p* = 0.004) and lower burst power (*r* = 0.87, *p* < 0.001).

**Figure 4.**
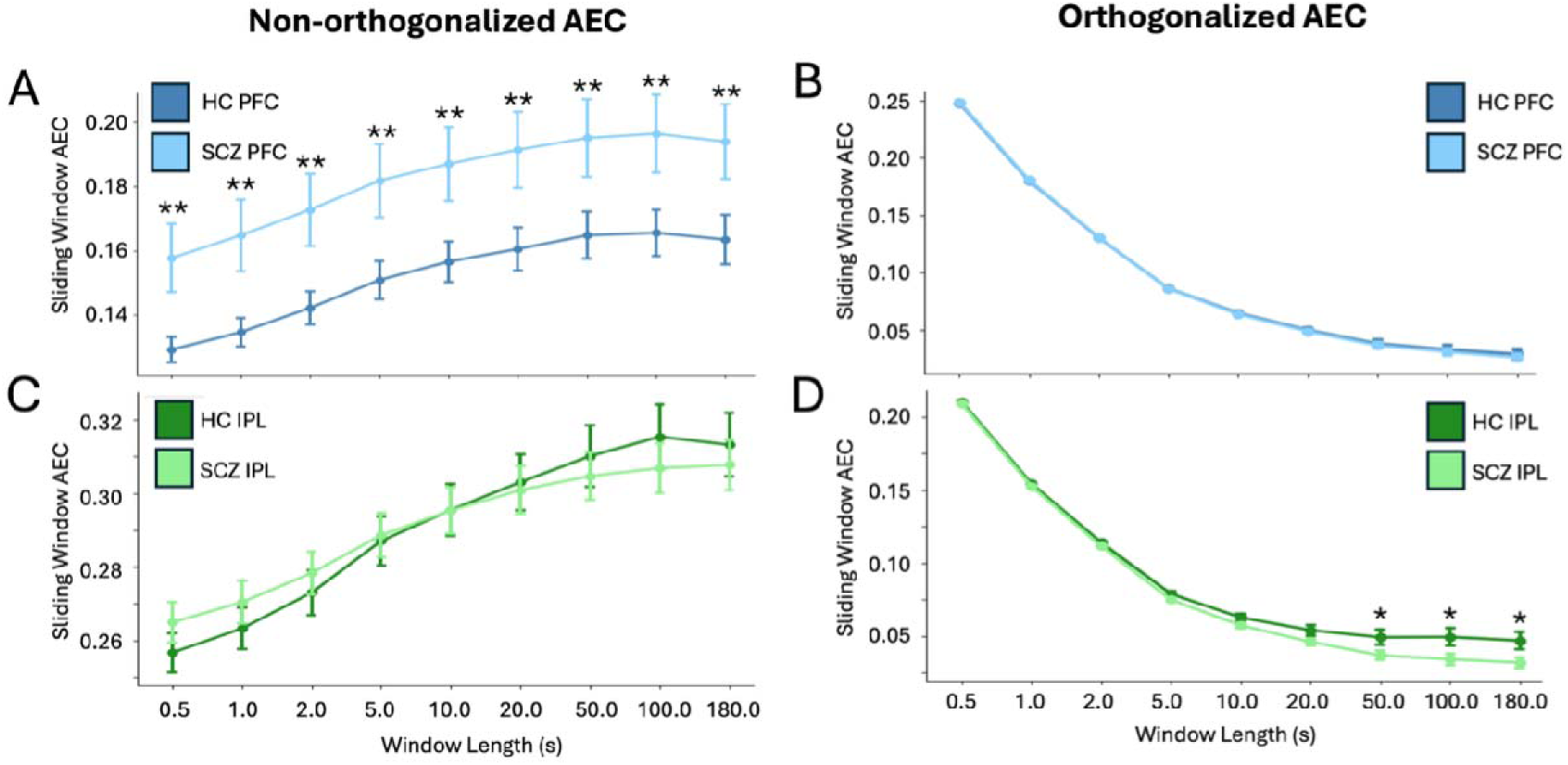
Sliding window AEC connectivity for IPL and PFC regions of interest. Mean non-orthogonalized (A) and orthogonalized (B) AEC values within PFC regions for sliding time windows of lengths between 0.5 seconds and 180 seconds (entire recording). AEC values are plotted for healthy controls (dark blue) and patients (light blue). Mean non-orthogonalized (C) and orthogonalized (D) AEC values within IPL regions. AEC values are plotted for healthy controls (dark green) and patients (light green). Error bars indicate standard error of the mean. Asterisks denote statistical significance (*: p<0.05; **: p<0.01).

Together, the AEC findings indicate that the increased PFC burst synchrony is a zero-lag phenomenon: present in non-orthogonalized AEC, absent after orthogonalization, consistent across all observation timescales, and not attributable to spatial proximity. Reduced IPL coupling is a delayed, slower timescale phenomenon: detectable only in orthogonalized AEC, emerging only at window lengths of 50 s or longer, and linked to shorter burst duration.

### 3.4 Burst-speech and symptom associations

Within patients (n = 23), the six group-discriminating burst features reduced to three principal components (PC1 36%, PC2 32%, PC3 16% of variance; cumulative 85%) (see Figure 5A). PC1 loaded on the A-P burst gradient; PC2 loaded on between- and within-prefrontal burst synchrony; and PC3 loaded predominantly on inferior parietal (IPL) burst duration (loading +0.81) with the A-P burst power gradient (+0.52). The speech organization component derived from syntactic and semantic complexity features (50% of variance) indexed lower clause count, syntax depth, and perplexity and greater consecutive similarity (see Figure 5B).

**Figure 5.**
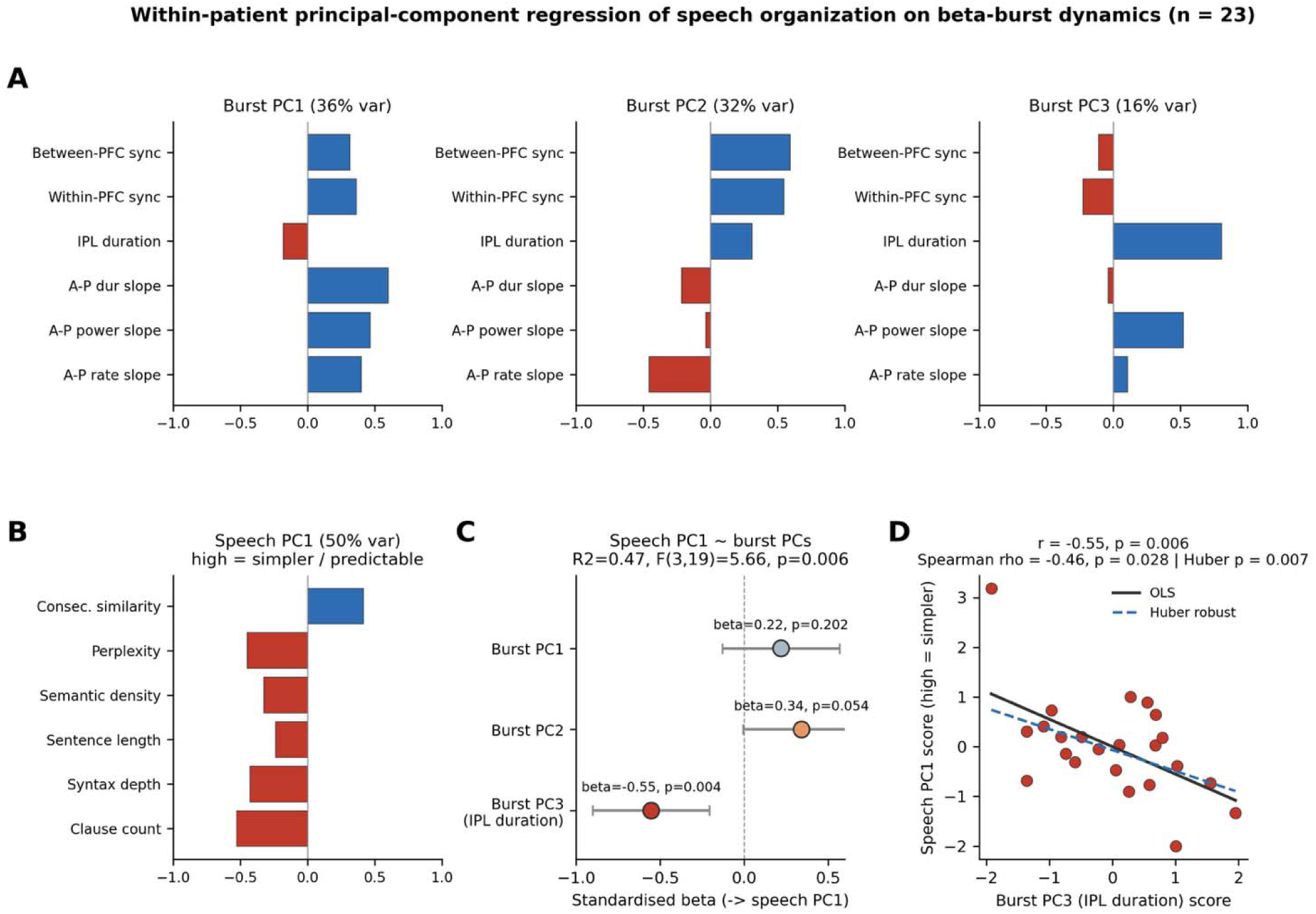
Within-patient principal-component regression of speech organization on beta-burst dynamics. The six beta-burst features showing significant group differences and the six semantic/syntactic speech features were each reduced by principal component analysis (z-scored within patients). **(A)** Loadings of the first three burst principal components: PC1 (36%) loads on the A-P burst gradients; PC2 (32%) on between- and within-prefrontal burst synchrony; and PC3 (16%) predominantly on inferior parietal (IPL) burst duration, with the A-P burst power gradient. **(B)** Loadings of the first speech component (speech PC1, 50% of speech variance), oriented so that higher scores denote simpler, more predictable speech (greater consecutive similarity; fewer clauses, lower syntax depth and perplexity). See speech feature description and group difference in Table S3. **(C)** Standardized coefficients (β, points; 95% CI, whiskers) from the multiple regression of speech PC1 on the three burst PCs (R² = 0.48, F(3,19) = 5.74, p = 0.006). The association was carried specifically by burst PC3/IPL burst duration (β = −0.55, p = 0.004); PC2 showed a weak trend (β = 0.34, p = 0.054) and PC1 no effect (β = 0.22, p = 0.202). Point colour denotes significance (red, p < 0.05; orange, p < 0.10; grey, n.s.). **(D)** Scatter of speech PC1 against burst PC3, with ordinary least-squares (solid) and Huber robust (dashed) fits. The association was significant and stable across outlier-resistant estimators (Pearson r = −0.55, p = 0.006; Spearman ρ = −0.46, p = 0.028; Huber robust regression p = 0.007), indicating it was not driven by a single influential case. Burst features: between-/within-PFC synchrony, IPL burst duration, and A-P burst duration/power/rate slopes; A–P, anteroposterior; PFC, prefrontal cortex; IPL, inferior parietal lobule.

A multiple regression of speech organization on the three burst PCs was significant (*R²* = 0.47, *adjusted R²* = 0.39; *F*(3,19) = 5.66, *p* = 0.006). The association was carried specifically by PC3 reflecting IPL burst duration (*β* = –0.55, *p* = 0.0036), with a weak contribution of prefrontal synchrony (PC2: *β* = 0.34, *p* = 0.054) but not the A-P burst gradient component (PC1: *β* = 0.22, *p* = 0.20) (see Figure 5C and 5D). Unpacking this at the single feature level, higher prefrontal synchrony was associated with shallow syntax structure (*r*=-0.53 to -0.61, *p*=0.002-0.009 for within and between PFC synchrony with clause count and syntax depth) while shorter IPL burst duration related to more predictable speech (*r*=-0.46, *p*=0.028 for consecutive similarity). Results were robust to outliers (Supplement 3.3, Table S5).

The same three burst PCs did not significantly predict clinical severity (clinical principal component; *R²* = 0.20, *F*(3,19) = 1.57, *p* = 0.23). A trend-level contribution was observed for global A-P burst component (PC1: *β* = –0.37, *p* = 0.092). Defined daily dose did not correlate with clinical severity (*r*(21) = 0.18, *p* = 0.42). Burst dynamics therefore showed partially dissociable correlates: IPL burst duration and to some extent prefrontal synchrony tracked speech organization, but not the overall symptom burden.

See partial least squares analysis of the full burst and speech feature sets with concordant burst-speech association in the Supplement (Supplementary 3.1-3.2, Figure S6-9).

## 4. Discussion

Beta abnormalities in schizophrenia are expressed through spatiotemporal changes in transient beta events rather than uniform changes in average beta power. Patients showed a cortex-wide reversal of the normal posterior-to-anterior gradient of burst rate, power, and duration, with two dissociable abnormalities within the DMN. Prefrontal regions showed excessive instantaneous co-occurrence of burst events. The inferior parietal lobule showed reduced burst duration and impaired slow-timescale coupling. Together, these features specifically predicted semantic and syntactic disorganization in spontaneous speech rather than the overall illness burden. These burst-level observations reconcile the heterogeneous prior resting-state reports of elevated beta power, altered network connectivity, and reduced posterior temporal stability of oscillatory activity in schizophrenia (Table S1). The findings are consistent with spatially and mechanistically distinct failures of predictive representations at anterior and posterior nodes of the cortical hierarchy in schizophrenia.

Increased burst synchrony within and between prefrontal DMN regions was mirrored by non-orthogonalized AEC and was present consistently across timescales. Synchrony differences increased with inter-regional Euclidean distance. This pattern identifies the prefrontal effect as an increase in instantaneous, zero-lag burst co-occurrence rather than an artifact of volume conduction or event detection. It is consistent with evidence that electrophysiological power-correlation networks reflect co-occurrence of transient high-amplitude events rather than sustained coupling ^24^, and with the view that beta bursts function as discrete computational events that stabilise and reinstate internal representations ^17,46^, Excessive instantaneous co-occurrence of prefrontal burst events may reflect over-stabilisation of internally generated representational states, providing a burst-level mechanism (i.e., increased coincidence) for prior reports of elevated frontal beta power and connectivity in schizophrenia ^47,48^. The contribution of prefrontal synchrony to speech syntax is consistent with the role of syntactic control on lexico-semantic choices and its impoverishment in schizophrenia ^49^. PFC over-synchrony may generate context-insensitive predictions that impoverish the representational input to the IPL.

Reduced burst duration in the IPL was accompanied by reductions in orthogonalized AEC at window lengths of 50 s or longer, which directly correlated with burst duration, with no corresponding reduction in non-orthogonalized AEC. This profile identifies the posterior effect as a deficit in sustained, time-delayed coupling that is a downstream consequence of burst brevity, providing a mechanistic explanation of previously reported reductions in long-range temporal correlations ^50–52^. Shorter bursts narrow the temporal window during which neural representations can be stabilized and broadcast across distributed regions ^20,21^. In the IPL, a hub for contextual grounding and semantic integration, sufficient burst duration may enable the maintenance of contextual representations across the timescales required for coherent language production; when this is impaired, word choices may be driven by higher consecutive similarity i.e., local rather than global context. These findings offer a burst-level account for previously reported reductions in long-range temporal correlations of beta oscillations in schizophrenia ^51,52^.

Complementary task-based evidence from the same cohort shows reduced event-related beta modulation in frontal regions during perception and motor action tasks ^53^. The combination of excessive anterior (PFC) resting synchrony, reduced posterior (IPL) burst duration and impaired distributed task-evoked modulation suggests context-insensitive top-down signalling: prefrontal predictions over-stabilised at rest and insufficiently reconfigured with contextual demands.

The cortex-wide group-by-position interaction for burst rate, power and duration indicates that the observed DMN findings are the focal expression of a broader reorganization of beta burst architecture across the entire cortex. Within the DMN, the gradient resolved to mechanistically distinct anterior and posterior effects: the prefrontal abnormality is a zero-lag, instantaneous phenomenon present at all timescales; the IPL abnormality is a delayed, slow-timescale phenomenon absent until zero-phase-lag components are removed. Within a predictive processing framework, these correspond to complementary failure modes: top-down predictions excessively instantiated at prefrontal sites while contextual evidence fails to be maintained long enough for posterior integration. Two interpretive caveats apply. First, the predictive processing account is a plausible mechanistic framework, not a demonstrated functional assignment; attentional suppression, arousal regulation, and network excitability fluctuations remain viable alternative accounts. Second, at the individual level, prefrontal synchrony and IPL burst duration were weakly negatively correlated (also see distinct loading on PCs in burst-speech regression). The two effects may reflect partially independent neurobiological processes affecting different subsets of patients rather than co-occurring consistently within individuals. Thus, the anterior-posterior burst imbalance should be treated as a group-level description rather than a demonstration of a mechanistic dissociation at an individual level.

## 5. Limitations

Several limitations apply. The modest sample restricts generalisability, and the burst-speech and burst-clinical associations are exploratory pending replication in larger cohorts. Sex was considered during recruitment by age- and sex-matching patients and controls, and sex distribution is reported in Table 1. Gender identity and socio-cultural gender-related variables were not systematically assessed; therefore, sex- or gender-specific effects could not be evaluated. MEG does not resolve laminar-specific signalling, limiting inference about whether prefrontal synchrony arises from superficial or deep cortical layers, as the predictive processing account would predict. Zero-phase-lag AEC retains some sensitivity to volume conduction; phase-based or laminar-resolved methods are the more stringent test. Beta burst metrics can be estimated reliably with 2-3 minute recordings ^54^, but the short epoch may increase variance in synchrony estimates. The eyes-open resting paradigm may suppress posterior beta via visual drive and requires replication with eyes-closed recordings. The cross-sectional design prevents causal inference, although associations with speech and clinical measures support functional relevance.

## 6. Conclusions

In summary, patients with schizophrenia exhibited a cortex-wide reversal of the anterior-posterior gradient of beta burst dynamics, with two mechanistically distinct abnormalities in the DMN: excessive instantaneous co-occurrence of prefrontal burst events, and reduced burst duration and slow-timescale coupling in the inferior parietal lobule. These findings suggest a hierarchical imbalance in which prefrontal predictions are over-stabilized while contextual representations in the parietal cortex are temporally too brief to sustain integration, both contributing to disorganization in spontaneous speech. By demonstrating that transient burst organization rather than spectral power captures these group differences, we report disrupted beta burst spatiotemporal architecture as a candidate mechanism linking intrinsic network dysfunction to thought disorder in schizophrenia.

## Acknowledgments

ChatGPT was used only to assist in identifying portions of the human-written manuscript that could be shortened. All AI-generated suggestions were critically reviewed, edited, and approved by the authors.

## Author contribution

**Hsi T. Wei:** Conceptualization, Software, Formal Analysis, Resources, Writing – Original Draft, Visualization, Project Administration. **Lindsey Power:** Software, Formal analysis, Writing - original draft, Writing - Review & Editing, Visualization. **Rukun Dou:** Software, Formal analysis. **Sylvain Baillet:** Conceptualization, Methodology, Writing – Review & Editing, Funding acquisition, Supervision. **Lena Palaniyappan:** Conceptualization, Methodology, Writing – Review & Editing, Funding acquisition, Supervision.

## Data Availability

Data generated or analyzed during this study are available from the corresponding author upon reasonable request.

## References

1. Rocca P, Galderisi S, Rossi A, et al. Disorganization and real-world functioning in schizophrenia: Results from the multicenter study of the Italian Network for Research on Psychoses. Schizophrenia Research 2018;201:105–12.

2. Ventura J, Thames AD, Wood RC, et al. Disorganization and Reality Distortion in Schizophrenia: A Meta-Analysis of the Relationship between Positive Symptoms and Neurocognitive Deficits. Schizophr Res 2010;121:1–14.

3. Huang H, Qin X, Xu R, et al. Default Mode Network, Disorganization, and Treatment-Resistant Schizophrenia. Schizophr Bull 2025;52:sbaf018.

4. Wold KF, Kreis IV, Åsbø G, et al. Long-term clinical recovery and treatment resistance in first-episode psychosis: a 10-year follow-up study. Schizophrenia (Heidelb) 2024;10:69.

5. Brown M, Kuperberg GR. A Hierarchical Generative Framework of Language Processing: Linking Language Perception, Interpretation, and Production Abnormalities in Schizophrenia. Front Hum Neurosci [Internet] Frontiers; 2015 [cited 2024 Sept 6];9. Available from: https://www.frontiersin.org/journals/human-neuroscience/articles/10.3389/fnhum.2015.00643/full

6. Palaniyappan L. Dissecting the neurobiology of linguistic disorganisation and impoverishment in schizophrenia. Semin Cell Dev Biol 2022;129:47–60.

7. Palaniyappan L, Venkatasubramanian G. The Bayesian brain and cooperative communication in schizophrenia. J Psychiatry Neurosci 2022;47:E48–54.

8. Chao ZC, Takaura K, Wang L, et al. Large-Scale Cortical Networks for Hierarchical Prediction and Prediction Error in the Primate Brain. Neuron 2018;100:1252–1266.e3.

9. Dimakou A, Pezzulo G, Zangrossi A, et al. The predictive nature of spontaneous brain activity across scales and species. Neuron 2025;113:1310–32.

10. Buckner RL, Andrews-Hanna JR, Schacter DL. The brain’s default network: anatomy, function, and relevance to disease. Ann N Y Acad Sci 2008;1124:1–38.

11. Northoff G, Heinzel A, de Greck M, et al. Self-referential processing in our brain—A meta-analysis of imaging studies on the self. NeuroImage 2006;31:440–57.

12. Smallwood J, Bernhardt BC, Leech R, et al. The default mode network in cognition: a topographical perspective. Nat Rev Neurosci 2021;22:503–13.

13. Andrews-Hanna JR, Reidler JS, Sepulcre J, et al. Functional-Anatomic Fractionation of the Brain’s Default Network. Neuron 2010;65:550–62.

14. Cao C, Liu W, Hou C, et al. Disrupted default mode network connectivity and its role in negative symptoms of schizophrenia. Psychiatry Research 2025;348:116489.

15. Pomarol-Clotet E, Salvador R, Sarró S, et al. Failure to deactivate in the prefrontal cortex in schizophrenia: dysfunction of the default mode network? Psychological Medicine 2008;38:1185– 93.

16. Whitfield-Gabrieli S, Thermenos HW, Milanovic S, et al. Hyperactivity and hyperconnectivity of the default network in schizophrenia and in first-degree relatives of persons with schizophrenia. Proceedings of the National Academy of Sciences Proceedings of the National Academy of Sciences; 2009;106:1279–84.

17. Spitzer B, Haegens S. Beyond the Status Quo: A Role for Beta Oscillations in Endogenous Content (Re)Activation. eNeuro [Internet] Society for Neuroscience; 2017 [cited 2024 Sept 5];4. Available from: https://www.eneuro.org/content/4/4/ENEURO.0170-17.2017

18. Vinck M, Uran C, Dowdall JR, et al. Large-scale interactions in predictive processing: oscillatory versus transient dynamics. Trends Cogn Sci 2025;29:133–48.

19. Feingold J, Gibson DJ, DePasquale B, et al. Bursts of beta oscillation differentiate postperformance activity in the striatum and motor cortex of monkeys performing movement tasks. Proceedings of the National Academy of Sciences Proceedings of the National Academy of Sciences; 2015;112:13687–92.

20. Lundqvist M, Rose J, Herman P, et al. Gamma and beta bursts underlie working memory. Neuron 2016;90:152–64.

21. Shin H, Law R, Tsutsui S, et al. The rate of transient beta frequency events predicts behavior across tasks and species. Kajikawa Y, editor. eLife eLife Sciences Publications, Ltd; 2017;6:e29086.

22. Rayson H, Moreau Q, Gailhard S, et al. Beta Burst Waveform Diversity: A Window onto Cortical Computation. Neuroscientist SAGE Publications Inc STM; 2025;10738584251390779.

23. Jones SR. When brain rhythms aren’t ‘rhythmic’: implication for their mechanisms and meaning. Current Opinion in Neurobiology 2016;40:72–80.

24. Hindriks R, Tewarie PKB. Dissociation between phase and power correlation networks in the human brain is driven by co-occurrent bursts. Commun Biol Nature Publishing Group; 2023;6:286.

25. Seedat Z, Quinn A, Vidaurre D, et al. The role of transient spectral ‘bursts’ in functional connectivity: A magnetoencephalography study. NeuroImage 2020;209:116537.

26. Briley PM, Liddle EB, Simmonite M, et al. Regional Brain Correlates of Beta Bursts in Health and Psychosis: A Concurrent Electroencephalography and Functional Magnetic Resonance Imaging Study. Biol Psychiatry Cogn Neurosci Neuroimaging 2021;6:1145–56.

27. Binder JR, Desai RH. The neurobiology of semantic memory. Trends Cogn Sci 2011;15:527–36.

28. Jefferies E. The neural basis of semantic cognition: converging evidence from neuropsychology, neuroimaging and TMS. Cortex 2013;49:611–25.

29. Zeng J, Liu Y, Zhao C, et al. Spatiotemporal Dynamics of Language Information Representation within the Left Inferior Parietal Lobule Network: An Intracranial SEEG Study. J Neurosci 2025;45:e0307252025.

30. Heilbron M, Armeni K, Schoffelen J-M, et al. A hierarchy of linguistic predictions during natural language comprehension. Proceedings of the National Academy of Sciences Proceedings of the National Academy of Sciences; 2022;119:e2201968119.

31. Hardy-Baylé M-C, Sarfati Y, Passerieux C. The cognitive basis of disorganization symptomatology in schizophrenia and its clinical correlates: toward a pathogenetic approach to disorganization. Schizophr Bull 2003;29:459–71.

32. Sharpe V, Mackinley M, Nour Eddine S, et al. Selective Insensitivity to Global Versus Local Linguistic Context in Speech Produced by Patients With Untreated Psychosis and Positive Thought Disorder. Biol Psychiatry 2025;S0006-3223(25)01243-0.

33. Cohen JD, Barch DM, Carter C, et al. Context-processing deficits in schizophrenia: converging evidence from three theoretically motivated cognitive tasks. J Abnorm Psychol 1999;108:120–33.

34. Matsumoto Y, Hayashi R, Takahashi H. Understanding semantic impairments in schizophrenia from a predictive coding perspective. Neurosci Res 2026;222:104990.

35. Liddle PF, Liddle EB. Imprecise Predictive Coding Is at the Core of Classical Schizophrenia. Front Hum Neurosci 2022;16:818711.

36. Lundqvist M, Miller EK, Nordmark J, et al. Beta: bursts of cognition. Trends in Cognitive Sciences Elsevier; 2024;28:662–76.

37. Dong D, Yao D, Wang Y, et al. Compressed sensorimotor-to-transmodal hierarchical organization in schizophrenia. Psychological Medicine 2023;53:771–84.

38. Holmes A, Levi PT, Chen Y-C, et al. Disruptions of Hierarchical Cortical Organization in Early Psychosis and Schizophrenia. Biological Psychiatry: Cognitive Neuroscience and Neuroimaging 2023;8:1240–50.

39. Karagoz AB, Moran EK, Barch DM, et al. Evidence for shallow cognitive maps in schizophrenia. Cogn Affect Behav Neurosci 2025;25:1192–209.

40. Alamia A, Gordillo D, Chkonia E, et al. Oscillatory Traveling Waves Provide Evidence for Predictive Coding Abnormalities in Schizophrenia. Biol Psychiatry 2025;98:167–74.

41. Andreasen NC. Thought, language, and communication disorders. I. Clinical assessment, definition of terms, and evaluation of their reliability. Arch Gen Psychiatry 1979;36:1315–21.

42. Kuperberg GR. Language in schizophrenia Part 1: an Introduction. Lang Linguist Compass 2010;4:576–89.

43. Cho B, Balles E, Mackinley M, et al. The DISCOURSE in psychosis (London Ontario): A speech dataset to examine communication disturbances in early-stage psychosis. Data Brief 2026;65:112517.

44. Schaefer A, Kong R, Gordon EM, et al. Local-Global Parcellation of the Human Cerebral Cortex from Intrinsic Functional Connectivity MRI. Cereb Cortex 2018;28:3095–114.

45. Colclough GL, Brookes MJ, Smith SM, et al. A symmetric multivariate leakage correction for MEG connectomes. Neuroimage 2015;117:439–48.

46. Sherman MA, Lee S, Law R, et al. Neural mechanisms of transient neocortical beta rhythms: Converging evidence from humans, computational modeling, monkeys, and mice. Proceedings of the National Academy of Sciences Proceedings of the National Academy of Sciences; 2016;113:E4885–94.

47. Canuet L, Ishii R, Pascual-Marqui RD, et al. Resting-State EEG Source Localization and Functional Connectivity in Schizophrenia-Like Psychosis of Epilepsy. PLoS One 2011;6:e27863.

48. Zeev-Wolf M, Levy J, Jahshan C, et al. MEG resting-state oscillations and their relationship to clinical symptoms in schizophrenia. Neuroimage Clin 2018;20:753–61.

49. Alonso-Sánchez MF, Hinzen W, He R, et al. Perplexity of utterances in untreated first-episode psychosis: an ultra–high field MRI dynamic causal modelling study of the semantic network. J Psychiatry Neurosci 2024;49:E252–62.

50. Alamian G, Pascarella A, Lajnef T, et al. Patient, interrupted: MEG oscillation dynamics reveal temporal dysconnectivity in schizophrenia. NeuroImage: Clinical 2020;28:102485.

51. Moran JK, Michail G, Heinz A, et al. Long-Range Temporal Correlations in Resting State Beta Oscillations are Reduced in Schizophrenia. Front Psychiatry 2019;10:517.

52. Nikulin VV, Jönsson EG, Brismar T. Attenuation of long-range temporal correlations in the amplitude dynamics of alpha and beta neuronal oscillations in patients with schizophrenia. NeuroImage 2012;61:162–9.

53. Wei HT, Boutet D, Dou R, et al. Distributed neurophysiological dynamics link perception, action, and language in schizophrenia [Internet]. bioRxiv; 2026 [cited 2026 Feb 23]. p. 2026.02.17.706352. Available from: https://www.biorxiv.org/content/10.64898/2026.02.17.706352v1

54. Pauls KAM, Nurmi P, Ala-Salomäki H, et al. Human sensorimotor resting state beta events and aperiodic activity show good test–retest reliability. Clinical Neurophysiology 2024;163:244–54.

