## Supplementary for "Dissociable Anterior and Posterior Beta-Burst Dynamics Track Thought Disorder in Schizophrenia"

**SI Text**

**1. Natural speech feature extraction**

Six speech features were extracted using NLP and LLM-based methods (Table S2). Semantic features captured how meaning unfolds across discourse and included consecutive sentence similarity, perplexity, and semantic density. Consecutive sentence similarity was computed as the average cosine similarity between sentence embeddings. Lexical predictability was indexed using perplexity. Semantic density was defined as the proportion of meaningful components within a sentence vector. Syntactic features indexed structural complexity based on part-of-speech parsing and included sentence length, syntactic tree depth, and clause count.

**2. Beta-burst detection and synchrony estimation**

Beta bursts were identified from time-frequency representations following established methods (main text ref. 21). The power threshold was selected to maximize the correlation between the percentage of suprathreshold TFR pixels and average beta power. For each burst, onset and offset times were defined by the full width at half maximum around the peak, corresponding to the last point before and first point after the peak where power was less than half of the peak value. Burst duration was defined as event offset minus onset, normalized peak power as the peak value expressed as a multiple of median power, and burst rate as the number of bursts per second. Inter-regional burst synchrony within the DMN was quantified by segmenting time series into 150 ms epochs centered on burst onset in region A and computing burst probability in region B using 50 ms sliding windows with a 10 ms step, yielding a temporal probability curve. Synchrony was defined as the probability of coincident bursting at time zero.

**3. Robustness analyses for burst-speech associations**

Rationale: Because the patient sample was modest (n = 23), the burst-speech findings could be vulnerable to three small-sample threats: confounding by diagnostic group when groups are pooled (Simpson’s paradox); optimisation/overfitting of multivariate cross-decomposition (PLS); and undue influence of individual cases. We therefore complemented the primary analysis with the robustness checks summarised below.

**3.1 Pooled versus within-group estimation**

The burst-speech PLS canonical correlation was significant in the pooled sample (r(46) = 0.56, permutation p < 0.001) (Figure S6). Because pooling two groups that differ on both burst and speech features can manufacture an association, we re-estimated the latent correlation within each group. The component was not significant in healthy controls (Figure S7: r(23) = 0.50, permutation p = 0.36) but was significant in patients (Figure S8: r(21) = 0.69, permutation p = 0.04; 120,000 permutations), indicating that the pooled effect was carried by the patient group and inflated by between-group separation. Permuting the burst block rather than the speech block gave equivalent results (patient permutation p = 0.04).

**3.2 Covariate adjustment**

Residualizing burst and speech features for diagnostic group reduced the pooled latent correlation (r = 0.45, permutation p = 0.06). Further adjustment rendered it non-significant: group + age + sex (Figure S9: r = 0.47, permutation p = 0.10) and group + age + sex + education (r = 0.47, permutation p = 0.07). The burst-speech latent correlation thus did not survive adjustment for demographic covariates in the combined sample, although nearing significance.

**3.3 Influential-case diagnostics and robust estimators**

One patient (MEG030) was statistically influential on the burst PC3-speech relationship (Cook’s distance = 1.05; leverage = 0.21). We therefore re-estimated the association with outlier-resistant methods (Table S5). It remained significant under rank correlation, biweight midcorrelation, and Huber and Tukey-biweight robust regression, and a Theil–Sen slope and bias-corrected bootstrap interval both excluded zero. Simple deletion of MEG030 left a trend (r = −0.40, p = 0.064), as expected from the loss of one observation under a non-robust estimator. At the single-feature level the most robust association was between inferior parietal burst duration and consecutive similarity (r = −0.46, p = 0.028), which was essentially unchanged after removing MEG030 (r = −0.44, p = 0.040).

**3.4 Specificity**

The same three burst PCs did not predict the clinical severity component (R² = 0.20, F(3,19) = 1.57, p = 0.23), and the inferior parietal component specifically was unrelated to severity (β = −0.06, p = 0.77). The speech association was therefore not a generic illness-severity effect; if anything, symptom severity related, at trend level, to a different (global anteroposterior/prefrontal-synchrony) burst dimension (PC1 β = −0.37, p = 0.09).

**3.5 Summary**

Across pooled, within-group, covariate-adjusted, dimension-reduced and outlier-resistant analyses, the burst–speech association was consistently (i) specific to patients, (ii) carried by inferior parietal burst duration, and (iii) robust to the single influential case under rank-based and robust estimators, while being attenuated by demographic covariate adjustment and only marginal in the multivariate PLS permutation test. Given the sample size, we interpret it as a reproducible but modest effect that warrants replication in a larger patient cohort.

**SI Tables**

**Table S1. Key studies on beta deviation in the DMN at rest in schizophrenia**

| **Study** | **Modality** | **Beta metric** | **Frontal beta** | **Temporal beta** | **Parietal / posterior beta** | **Main interpretation** | **Symptom association** |
| --- | --- | --- | --- | --- | --- | --- | --- |
| (32) | EEG | Spectral beta power | ↑ increased | ↑ increased | — | Replicated fronto-temporal beta elevation with posterior reductions | Linked to subclinical traits associated with vulnerability to psychosis |
| (33) | EEG-LORETA | Beta connectivity | ↑ increased connectivity | ↑ increased | — | Temporo-prefrontal hyperconnectivity | Correlated with positive symptoms |
| (34) | EEG | Beta long-range temporal correlations | — | — | ↓ reduced LRTC posterior | Reduced temporal stability of beta oscillations | Network instability hypothesis |
| (35) | EEG | Group-ICA derived slow beta / fast beta spectral components | ↑ increased fronto-central slow beta in SZ | — | — | Suggests fronto-central slow beta elevation and possible endophenotypic / excitability-related abnormality | No beta correlation with PANSS positive, negative, or total |
| (36) | MEG | Source beta power | ↑ increased | ↑ increased | — | Left-hemisphere beta elevation | Associated with negative symptoms |
| (37) | EEG | Beta long-range temporal correlation (LRTC) | — | — | ↓ reduced posterior LRTC | Beta temporal organization impaired | Not symptom-specific |
| (38) | EEG | Beta network connectivity | ↑ frontal network | — | — | Abnormal global oscillatory networks | PANSS correlations |
| (39) | EEG | Beta network connectivity | ↑ stronger frontal-parietal connectivity | — | — | Abnormal network efficiency | Correlated with PANSS scores |

**Table S2. NLP/LLM models for speech feature extraction**

| Feature | Feature domain | Model / parser used | Implementation details | English/French handling |
| --- | --- | --- | --- | --- |
| Consecutive sentence similarity | Semantic coherence | intfloat/multilingual-e5-base | Sentences were tokenized with the corresponding HuggingFace tokenizer. Sentence-level embeddings were extracted from the first-token representation of the final hidden layer. Consecutive similarity was computed as the cosine similarity between adjacent sentence embeddings and averaged across the transcript. | This embedding model is multilingual and was applied to both English and French transcripts. |
| Perplexity | Lexical / sequential predictability | ibm-granite/granite-3.3-2b-base | The full transcript was tokenized with the corresponding HuggingFace tokenizer and passed to a decoder-only causal language model. Perplexity was computed as the exponentiated cross-entropy loss over next-token prediction. | The same decoder-only model was applied to English and French transcripts. Because the model was not used as a language-specific normative benchmark, perplexity values were interpreted comparatively within the study rather than as absolute language-specific estimates. |
| Semantic density | Semantic richness / compactness | Vocabulary-restricted version* of the pre-trained Google News Word2Vec model (word2vec-google-news-300), implemented using Gensim | Content words were selected using POS tags, lemmatized, lowercased, and retained only if present in the Word2Vec vocabulary. Sentence-level semantic vectors were calculated by averaging retained word vectors. A gradient-based reconstruction procedure estimated the number of meaning-bearing components required to approximate each sentence vector; semantic density was calculated as the number of selected meaning components divided by the number of retained content words. | The same Gensim Word2Vec model was applied to English and French transcripts. Because Gensim is the implementation rather than a language-specific model family, bilingual coverage depends on the vocabulary of the specific Word2Vec model; therefore, only lemmatized content words present in the model vocabulary were retained. |
| Sentence length | Syntactic production quantity | stanza | Sentences were parsed using stanza.Pipeline(lang, processors="tokenize,mwt,pos,lemma"). Sentence length was calculated as the number of parsed tokens per sentence and averaged across the transcript. | Each subject consistently used English or French in the speech task. Transcript language was assigned at the file level: files containing “FR” were processed as French (lang="fr"), and all other files were processed as English (lang="en"). English analyses used Penn-style xpos tags; French analyses used Universal POS upos tags. |
| Syntax depth | Syntactic complexity | stanza + nltk.RegexpParser | POS-tagged and lemmatized tokens were passed to nltk.RegexpParser using language-specific grammar rules. Syntax depth was calculated as the height of each resulting chunk tree and averaged across sentences. | Separate English and French grammar rules were used. English parsing used Penn-style xpos tags; French parsing used Universal POS upos tags. |
| Clause count | Syntactic complexity | stanza + nltk.RegexpParser | Clause count was calculated as the number of chunk-tree subtrees labelled CLAUSE in each sentence and averaged across sentences. | Separate English and French chunking rules were used. English parsing used Penn-style xpos tags; French parsing used Universal POS upos tags. |

*****Word embeddings were derived from a vocabulary-restricted version of the pre-trained Google News Word2Vec model (word2vec-google-news-300), implemented using Gensim. The original model contains 300-dimensional vectors trained on the Google News corpus. To reduce the vocabulary for semantic density estimation, the original model was filtered to retain only common English and French words. Specifically, lists of the 50,000 most frequent English and French words were downloaded from the hermitdave/FrequencyWords repository; the top 15,000 words from each language were lemmatized using Stanza, lowercased, deduplicated, and used as a candidate vocabulary. Only candidate words present in the original Google News Word2Vec model were retained. The resulting reduced model was saved as a Gensim KeyedVectors object and contained 300-dimensional vectors for 15,855 vocabulary items. This vocabulary restriction was applied to improve semantic density estimation by avoiding optimization over the full Google News vocabulary of approximately 3 million tokens, which included many irrelevant or noisy entries. The reduced model was saved using Gensim 4.3.3 and Python 3.12.9.

**Table S3. Speech feature group difference**

| **Speech aspect** | **Speech feature** | **Feature Description** | **Group difference (PT - HC)** |
| --- | --- | --- | --- |
| Semantics | Consecutive similarity | Sentence-level cosine similarity. Higher similarity score indicates greater similarity between neighboring sentences in the embedding space. | *t*(45.79) = 3.91, *p* = 0.00030 |
|  | Perplexity | Perplexity of a generative model when evaluating the text from a participant. Higher perplexity reflects a lower capacity of the model to predict the subject’s word choice. | *t*(45.28) = -2.03, *p* = 0.048 |
|  | Semantic density | Ratio of meaningful to content words in a sentence, where meaningful components are derived via vector unpacking, applying gradient descent to word2vec sentence embeddings to approximate the original semantic vector. A higher value shows higher semantic density. | *t*(45.22) = .67, *p* = .51 |
| Syntactic | Sentence length | Number of words in a sentence. | *t*(43.86) = -4.42, *p* = 0.000064 |
|  | Syntax depth | Depth of the syntax tree obtained by parsing a sentence through part-of-speech tagging. A higher value indicates a complex syntax. | *t*(25.47) = -2.72, *p* = 0.012 |
|  | Clause count | Number of clauses as recognized by pattern matching on the part-of-speech tags. | *t*(45.66) = -3.70, *p* = 0.00058 |

**Note:** The 7 features index two complementary aspects of spontaneous discourse: *semantic* organization (how meaning unfolds and how predictable word choices are) and *syntactic* organization (how structurally elaborated each utterance is). Together, the patient profile is one of simpler, shorter, more syntactically shallow and more repetitive speech with more predictable lexical choices i.e. reduced semantic diversity and syntactic complexity, which is consistent with linguistic disorganization and impoverishment of spontaneous discourse. Higher scores on consecutive similarity and lower scores on perplexity, sentence length, syntax depth and clause count all denote this simpler/less-elaborated pole.

**Table S4. Clinical measures and their loadings on the clinical PC**

| **Clinical scores** | **Clinical measure** | **1st component loadings** |
| --- | --- | --- |
| SOFAS | **Social and Occupational Functioning Assessment Scale** | **-.34** |
| PANSS-P1 | **Delusion** | **.36** |
| PANSS-P2 | Conceptual disorganization | .26 |
| PANSS-P3 | **Hallucinatory Behavior** | **.30** |
| PANSS-N1 | **Blunted affect** | **.33** |
| PANSS-N4 | **Passive/apathetic social withdrawal** | **.37** |
| PANSS-N6 | Lack of spontaneity and flow of conversation | .17 |
| PANSS-G5 | Mannerisms/Posturing | .17 |
| PANSS-G9 | Unusual thought content | **.36** |
| CGI-S | Considering your total clinical experience with patients with schizophrenia, how severely ill has the patient been during the course of illness? | **.40** |

Note: Feature |loadings| ≥ .30 contribution to the component is bolded. SOFAS: The higher the score, the better the social functioning. PANSS and CGI-S: the higher the score, the worse the symptom.

**Table S5. Robustness of the inferior-parietal-burst-duration (burst PC3) - speech PC1 association in patients (n = 23)**

| **Estimator** | **Coefficient** | **p / 95% CI** |
| --- | --- | --- |
| Pearson correlation (reference) | r = −0.553 | p = 0.006 |
| Spearman rank correlation | ρ = −0.458 | p = 0.028 |
| Huber robust regression | slope = −0.422 | p = 0.007 |
| Bootstrap Pearson (10,000) | r = −0.553 | 95% CI [−0.779, −0.084] |

**SI Figures**

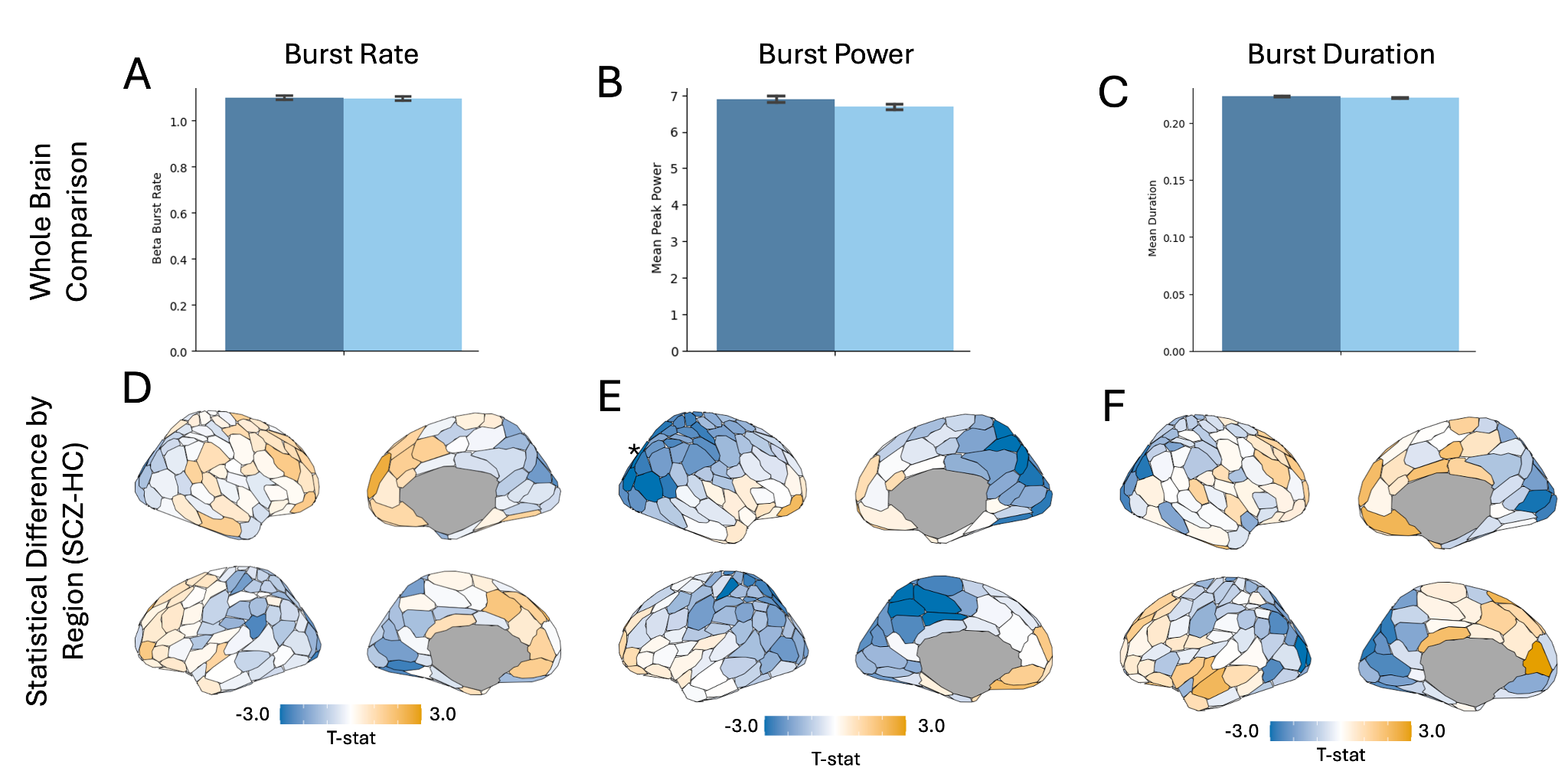

**Figure S1. Whole-brain beta burst characteristic summary.** (A-C) Mean burst rate (A), burst power (B), and burst duration (C) for healthy controls (dark blue) and patients (light blue) across subjects and brain regions. Error bars indicate standard error of the mean. No global differences in burst characteristics were observed. (D-F) Cortical maps showing the statistical difference (T-statistic) in burst rate (D), burst power (E), and burst duration (F) between patients and healthy controls for each Schaefer-200 region. Negative values (blue) indicate reduced values in SCZ compared to HC and positive values (orange) indicate increased values in SCZ compared to HC. Only one region was statistically significant for burst power: RH_DorsAttnA_SPL_1: T=4.59, FDR-p=0.00692. No significant effects were found for burst rate or duration.

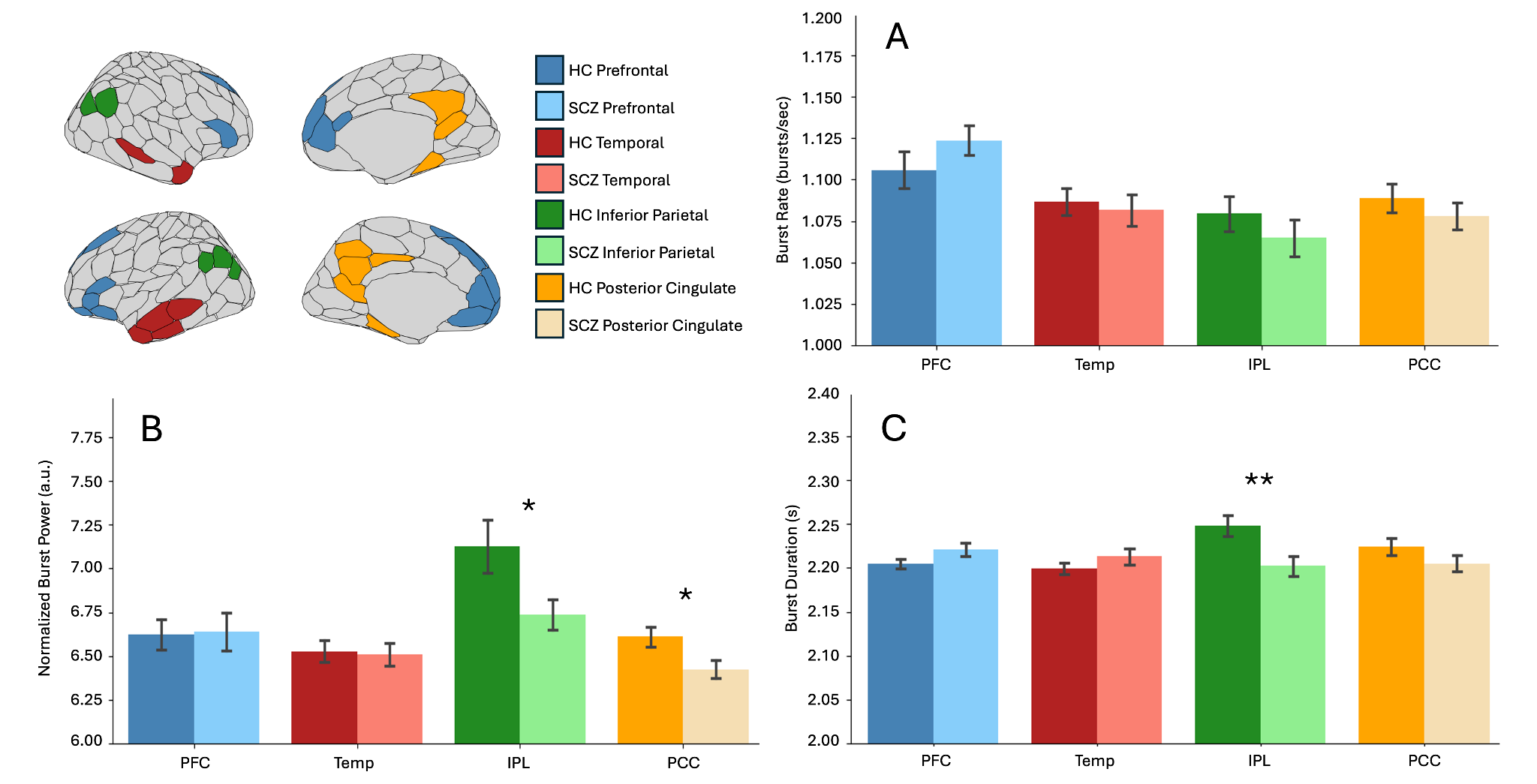

**Figure S2. DMN beta burst characteristics in schizophrenia and healthy controls.** Mean burst rate (A), burst power (B) and burst duration (C) for four DMN sub-regions: PFC (blue), Temporal (red), IPL (green), and PCC (orange). Error bars indicate standard error of the mean. Asterisks denote statistical significance (*: p<0.05, **: FDR-p<0.05).

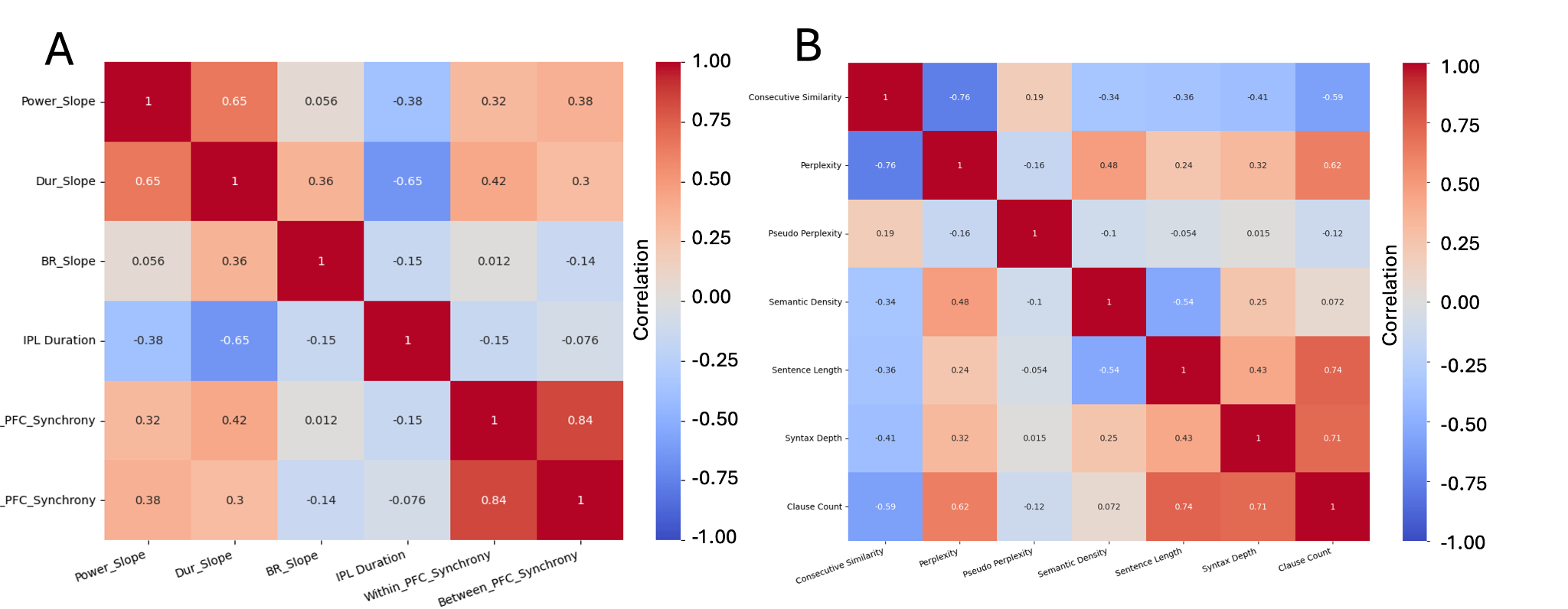
**Figure S3. Feature collinearity matrices for relevant burst and speech features.** (A) Matrix of Pearson correlation coefficients between each pair of significant burst features: A-P burst power slope, A-P burst duration slope, A-P burst rate slope, IPL burst duration, within left hemisphere PFC synchrony, and between hemispheres PFC synchrony. (B) Matrix of Pearson correlation coefficients between each pair of speech features: consecutive similarity, perplexity, pseudo perplexity, semantic density, sentence length, syntax depth, and clause count.

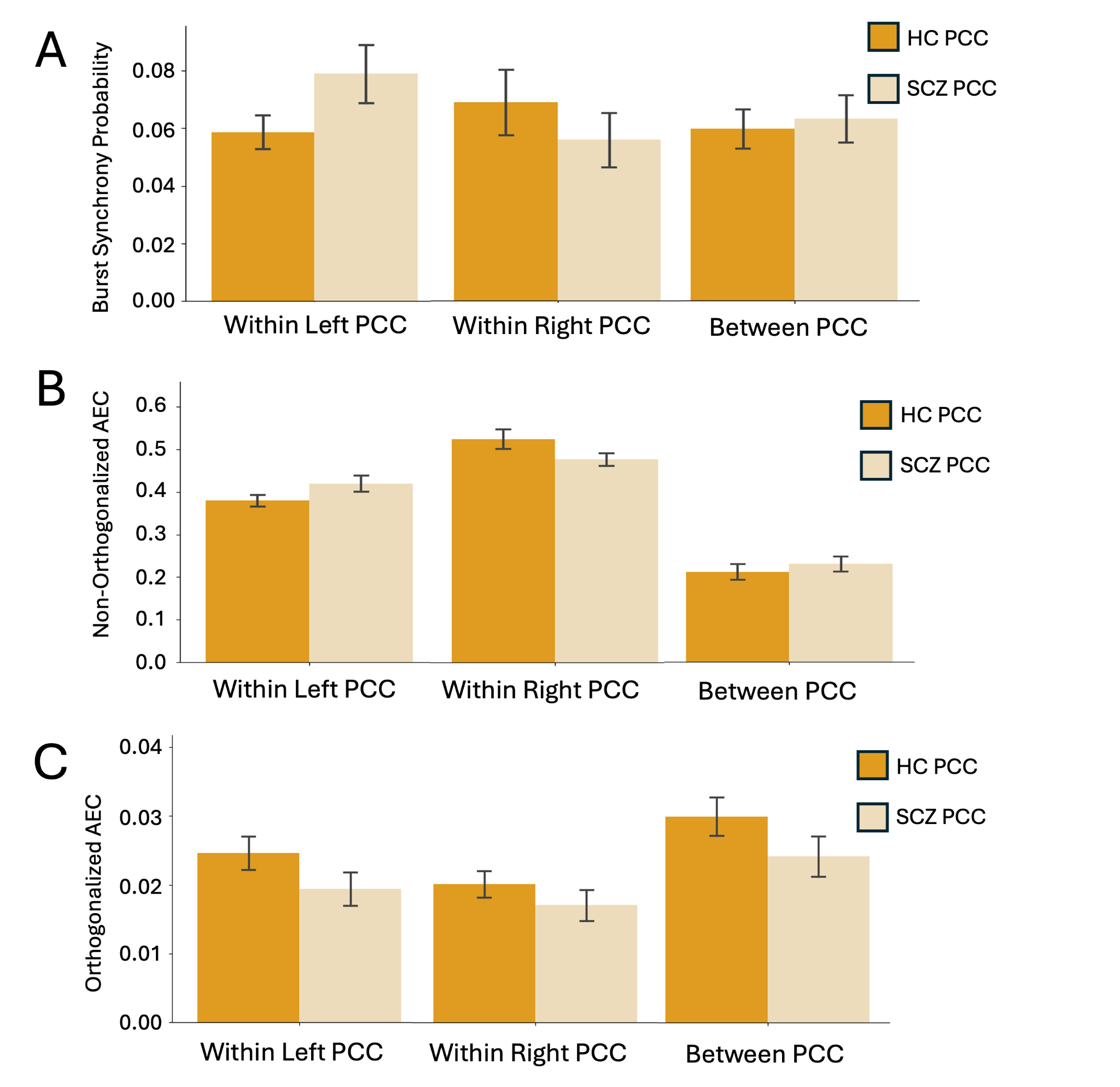

**Figure S4. Burst synchrony and connectivity metrics for Temporal and PCC regions of interest.** Mean values and standard error (error bars) are shown for burst synchrony (A-B), non-orthogonalized AEC (C-D), orthogonalized AEC (E-F), coherence (G-H), and imaginary coherence (I-J). The left column shows values within left temporal (left), within right temporal (center), and between left and right temporal (right) regions for healthy controls (dark red) and patients (light red). The right column shows values within left PCC (left), within right PCC (center) and between left and right PCC (right) for healthy controls (dark orange) and patients (light orange). Asterisks denote statistical significance (*: p<0.05, **: FDR-p<0.05).

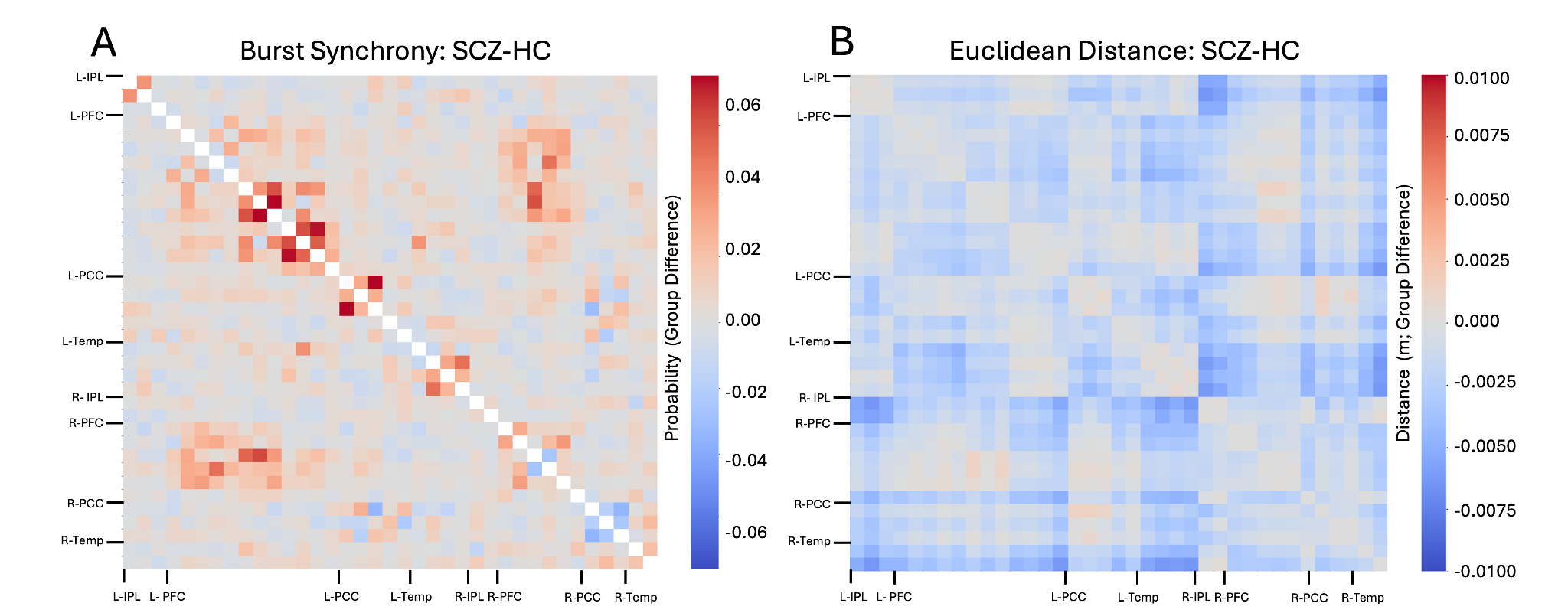
**Figure S5. Group difference matrices for burst synchrony and Euclidean distance.** (A) Group difference in synchrony (SCZ-HC) for each pair of regions in the DMN. (B) Group difference in distance between average region coordinates for each pair of regions in the DMN.

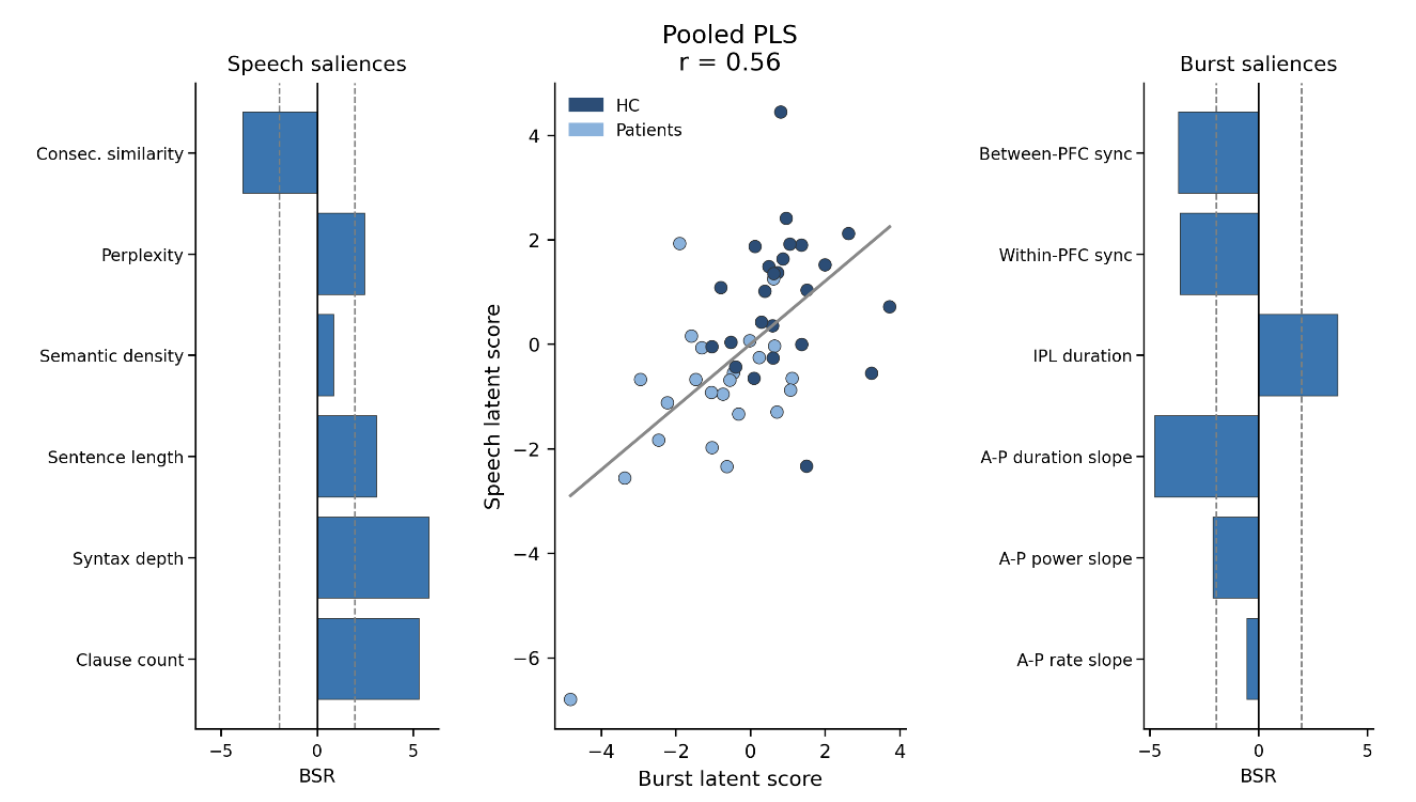

**Figure S6. Burst-speech PLSC on pooled participants**

Middle: Scatter plot of the latent burst (x-axis) and speech (y-axis) feature scores derived from the PLS canonical model. Healthy controls are coloured in dark blue and patients are coloured in light blue. Left: Each speech feature’s bootstrap ratio. Right: Each burst feature’s bootstrap ratio.

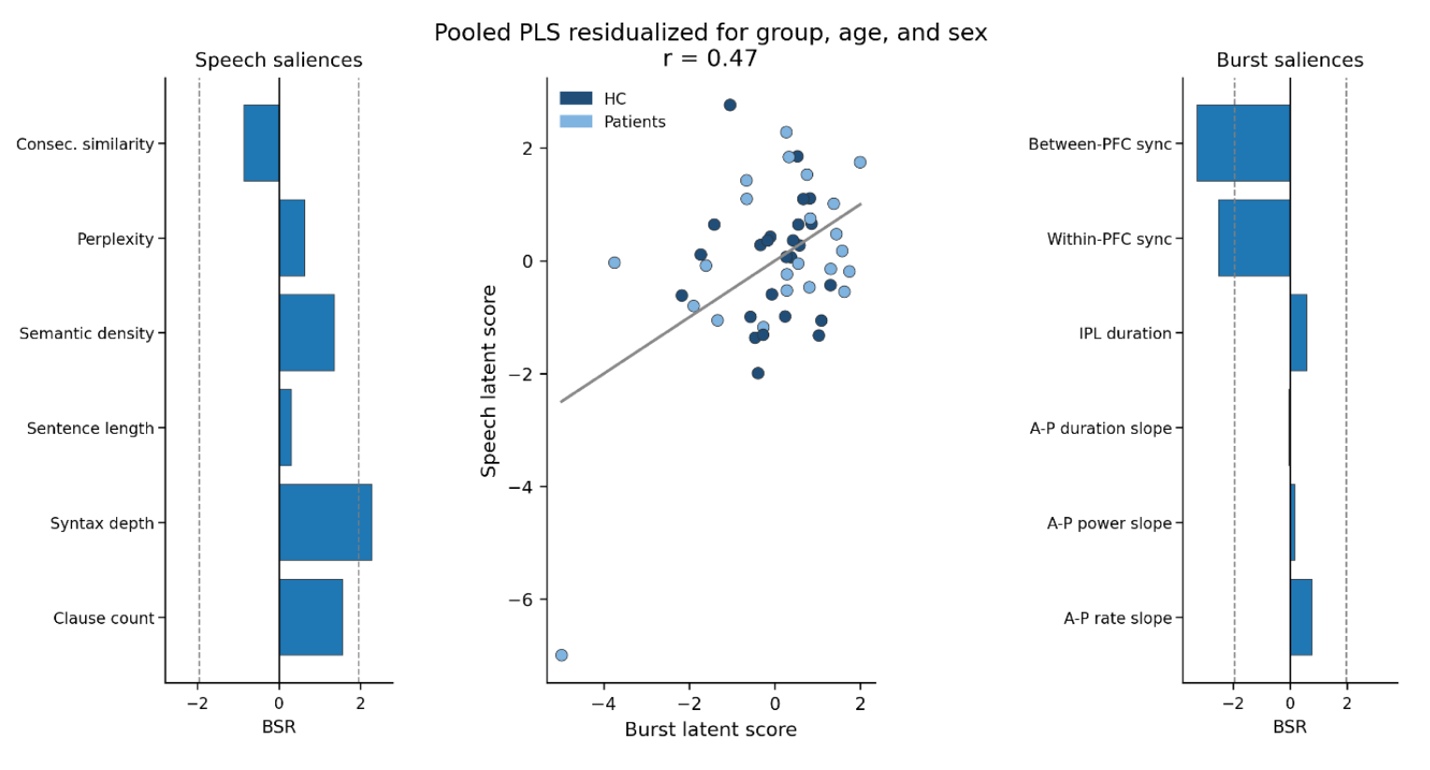

**Figure S7. Burst-speech PLSC on pooled participants after residualizing for group, age, and sex**

Middle: Scatter plot of the latent burst (x-axis) and speech (y-axis) feature scores derived from the PLS canonical model. Healthy controls are coloured in dark blue and patients are coloured in light blue. Left: Each speech feature’s bootstrap ratio. Right: Each burst feature’s bootstrap ratio.

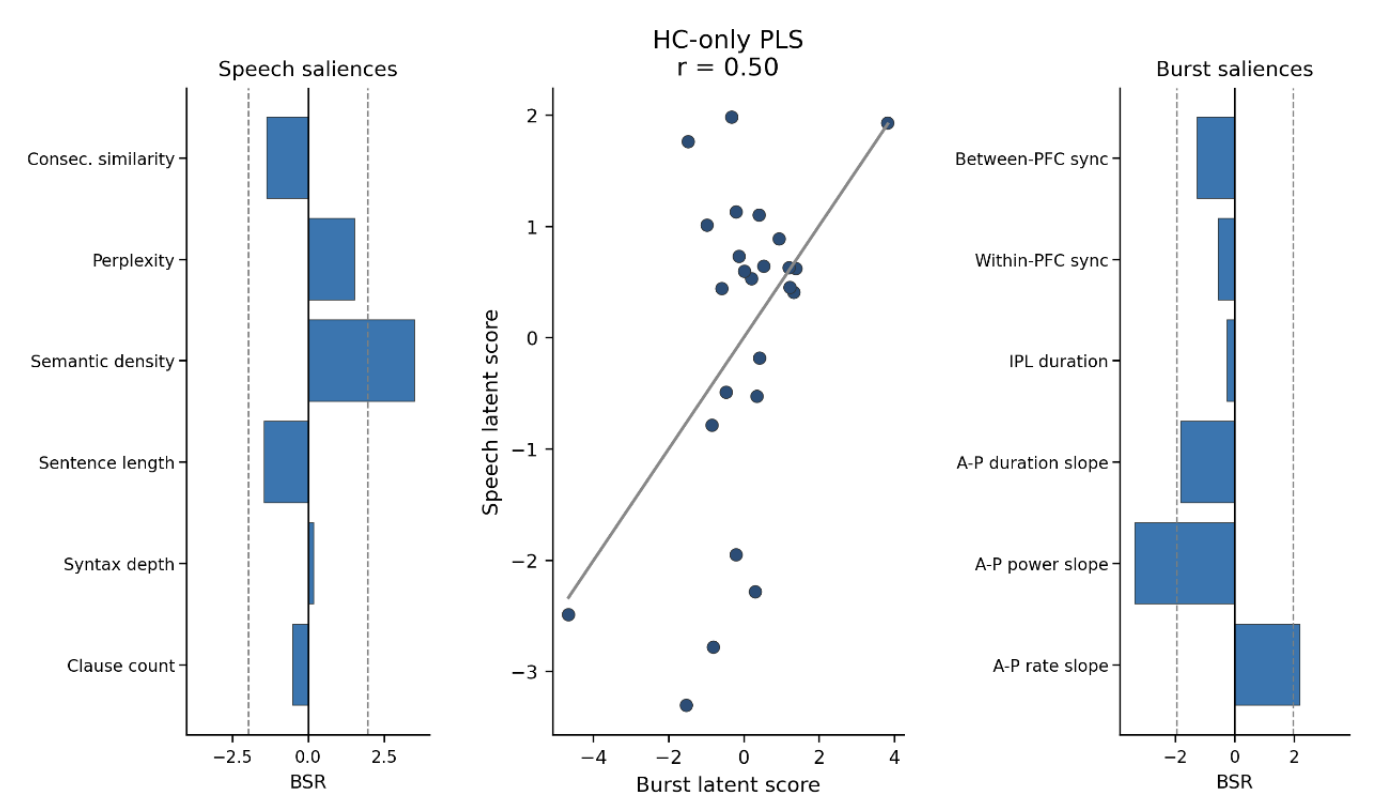

**Figure S8. Burst-speech PLSC on controls only**

Middle: Scatter plot of the latent burst (x-axis) and speech (y-axis) feature scores derived from the PLS canonical model. Left: Each speech feature’s bootstrap ratio. Right: Each burst feature’s bootstrap ratio.

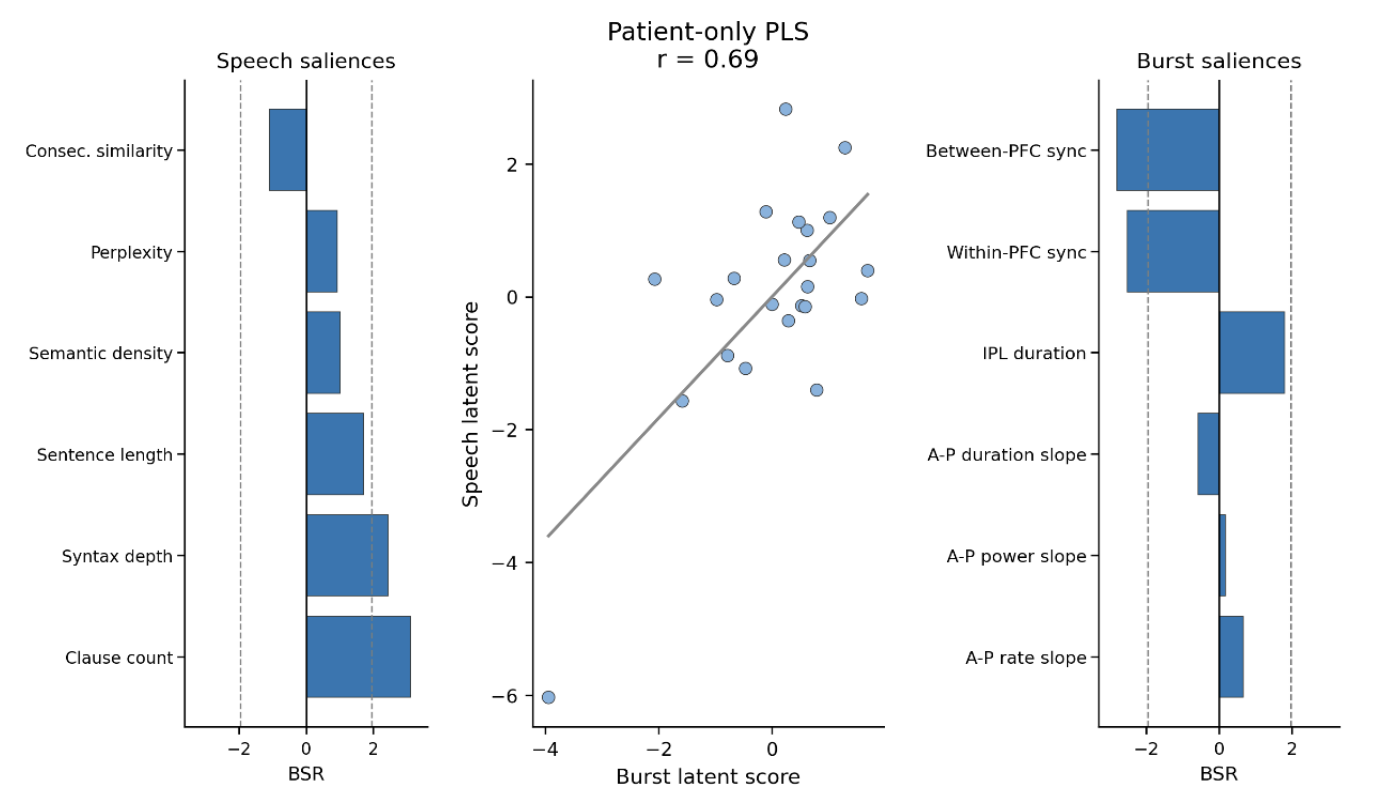

**Figure S9. Burst-speech PLSC on patients only**

Middle: Scatter plot of the latent burst (x-axis) and speech (y-axis) feature scores derived from the PLS canonical model. Left: Each speech feature’s bootstrap ratio. Right: Each burst feature’s bootstrap ratio.
